# Human skin fibrosis with iPSC-derived organoids reveals RUNX2-mediated fibroblast reprogramming

**DOI:** 10.64898/2026.09.14.751420

**Authors:** Jasim Khan, Sarmad Mehmood, Fuad Al Abir, Miao Wei, Suhail Muzaffar, Suha Mohiuddin, Shanrun Liu, Mohammad Waseem, Juheb Akhter, Zhipeng Liu, David K. Crossman, Masakazu Kamata, Jack Y. Chen, Chander Raman, Craig Elmets, Sixue Zhang, Mohammad Athar, Lin Jin

**Author notes:** These authors contributed equally (Jasim Khan, Sarmad Mehmood). Correspondence (S.Z), (M.A), (L.J).

## Abstract

Fibrotic skin diseases are characterized by persistent fibroblast activation and extracellular matrix remodeling, yet the mechanisms governing fibroblast state transitions remain incompletely understood. Here, we established a human iPSC-derived skin organoid model of fibrosis through chronic TGF-β stimulation. Single-cell RNA sequencing combined with immunofluorescence-based spatial analysis revealed dynamic fibroblast state transitions, spatial reorganization, and expansion of activated fibroblast populations during fibrotic remodeling. Integration with human scleroderma single-cell datasets demonstrated conserved fibroblast states and transcriptional programs between organoids and patient tissues. We further identified broad induction of RUNX2 in the dermal compartment during fibrosis, and RUNX2 depletion attenuated fibrotic marker expression. CUT&RUN profiling revealed RUNX2 occupancy at fibrosis-associated loci, including *RUNX1* and *LOXL2*. Using a machine learning-guided screening approach, we identified F0565-0303, a small molecule that suppressed RUNX2-dependent fibrotic programs in vitro and reduced fibrosis in a bleomycin-induced mouse model. Together, these findings establish human skin organoids as a platform for modeling fibrosis and nominate RUNX2 as a potential therapeutic target.

**One Sentence Summary:** Human skin organoids recapitulate fibrotic remodeling and reveal RUNX2 as a therapeutic target

## Introduction

Fibrotic remodeling of the skin is a hallmark of pathological conditions, including scleroderma, keloids and hypertrophic scars, radiation-induced fibrosis, chronic wounds, and graft-versus-host disease. It reflects dysregulated wound healing characterized by persistent fibroblast activation and excessive extracellular matrix (ECM) deposition. Central to this process is the emergence of activated fibroblast states that drive matrix production, tissue stiffening, and altered intercellular signaling ^(1–3)^. Transforming growth factor–β (TGF-β) signaling is a well-established driver of these processes, promoting myofibroblast differentiation and transcriptional reprogramming of fibroblasts toward profibrotic phenotypes ^(1,4–5)^. Despite advances in understanding key signaling pathways, the dynamic state transitions underlying fibrotic remodeling remain incompletely defined, particularly in human systems.

Single-cell transcriptomic studies have revealed substantial heterogeneity within dermal fibroblast populations, including matrix-producing, contractile, inflammatory, progenitor-like, and signaling-associated states ^(6–8)^. These findings suggest that fibroblast response to tissue injury and fibrosis may involve coordinated changes across multiple cellular states rather than a uniform activation program. In fibrotic diseases such as systemic scleroderma and keloid, these fibroblast populations undergo extensive remodeling, characterized by expansion of cells expressing ECM and profibrotic genes (e.g., *POSTN*, *COL11A1*, and *FN1*) and activation of TGF-β–responsive programs (e.g., *SERPINE1* and *CTGF*), along with increased expression of regulators associated with fibroblast activation and inflammation (e.g., *THY1* and *FAP*) ^(9–11)^. Complementary insights from mouse lineage-tracing and single-cell studies have further demonstrated dynamic fibroblast activation and state transitions during wound healing and fibrosis ^(12–14)^. However, most current understanding relies on static human biopsies or mouse models, limiting the ability to resolve human-specific temporal trajectories and regulatory programs that drive fibrotic progression. In parallel, widely used *in vitro* systems, including two-dimensional fibroblast cultures and “scar-in-a-jar” assays—which rely on exogenous collagen addition or macromolecular crowding to accelerate ECM deposition—lack three-dimensional tissue architecture, multicellular complexity, and physiological temporal progression, thereby constraining their ability to model fibroblast state transitions and fibrotic remodeling *in vivo* ^(5, 15–16)^. To overcome these limitations, we leveraged established human induced pluripotent stem cell (iPSC)-derived skin organoid platforms ^(17)^ and induced fibrotic remodeling using TGF-β, enabling analysis of fibroblast state transitions within a three-dimensional skin-like tissue architecture.

The Runt-related transcription factor 2 (RUNX2), best known for its role in osteogenic differentiation, has also been shown to contribute to fibroblast activation and tissue remodeling ^(18–21)^. Consistent with this broader profibrotic role, RUNX2 has also been additionally been linked with stromal activation and ECM remodeling in tumor-associated contexts ^(22–24)^. At the genomic level, RUNX2 binds promoters and enhancers to regulate lineage-specific transcriptional programs, including osteogenic and ECM–associated genes ^(25,26)^. However, its genome-wide targets and roles in fibroblast state transitions remain poorly defined.

Here, we establish a TGF-β–induced fibrotic model in iPSC-derived skin organoids to investigate fibroblast state dynamics and their regulatory mechanisms. Integrating single-cell transcriptomics and trajectory analysis, we define fibroblast populations, characterize their transitions, and delineate transcriptional reprogramming during fibrotic progression. We further compare these states with published single-cell datasets from scleroderma to assess disease relevance. We identify RUNX2 as a key regulator of fibrotic remodeling and delineate its genomic target modules using CUT&RUN sequencing. Finally, we demonstrate that pharmacological targeting of RUNX2 signaling attenuates fibrotic phenotypes, highlighting a potential therapeutic strategy.

## Results

### Time-resolved TGF-β stimulation induces fibrotic remodeling in human iPSC-derived skin organoids

To establish a human iPSC-derived platform for modeling dermal fibrosis, we generated skin organoids and matured them for 80 days using a previously published differentiation protocol ^(17)^. The resulting organoids developed distinct epidermal and dermal-like compartments that recapitulate key features of human skin architecture. Cell identities and lineage specification were validated by established lineage markers, including KRT14 and KRT10 for basal and suprabasal keratinocytes, respectively, PDGFRA for dermal fibroblasts, and SOX2 for hair follicle–associated dermal condensate cells (Fig. 1A). Given the central role of TGF-β signaling in fibroblast activation and ECM deposition across fibrotic diseases, we used TGF-β stimulation to model fibrotic remodeling in skin organoids ^(5)^. Organoids were exposed to TGF-β (10 ng/mL) every other day for up to 14 days, with untreated organoids serving as controls. To account for developmental stage-dependent effects, stimulation was initiated either at day 0 (14-day exposure) or day 6 (8-day exposure), and all organoids were harvested at a common endpoint on day 14 (Fig. 1A). Cell identities and lineage specification were validated using established markers, including KRT14 and KRT10 for basal and suprabasal keratinocytes, respectively, PDGFRA for dermal fibroblasts, and SOX2 for hair follicle-associated dermal condensate cells (Fig. 1A).

**Figure 1.**
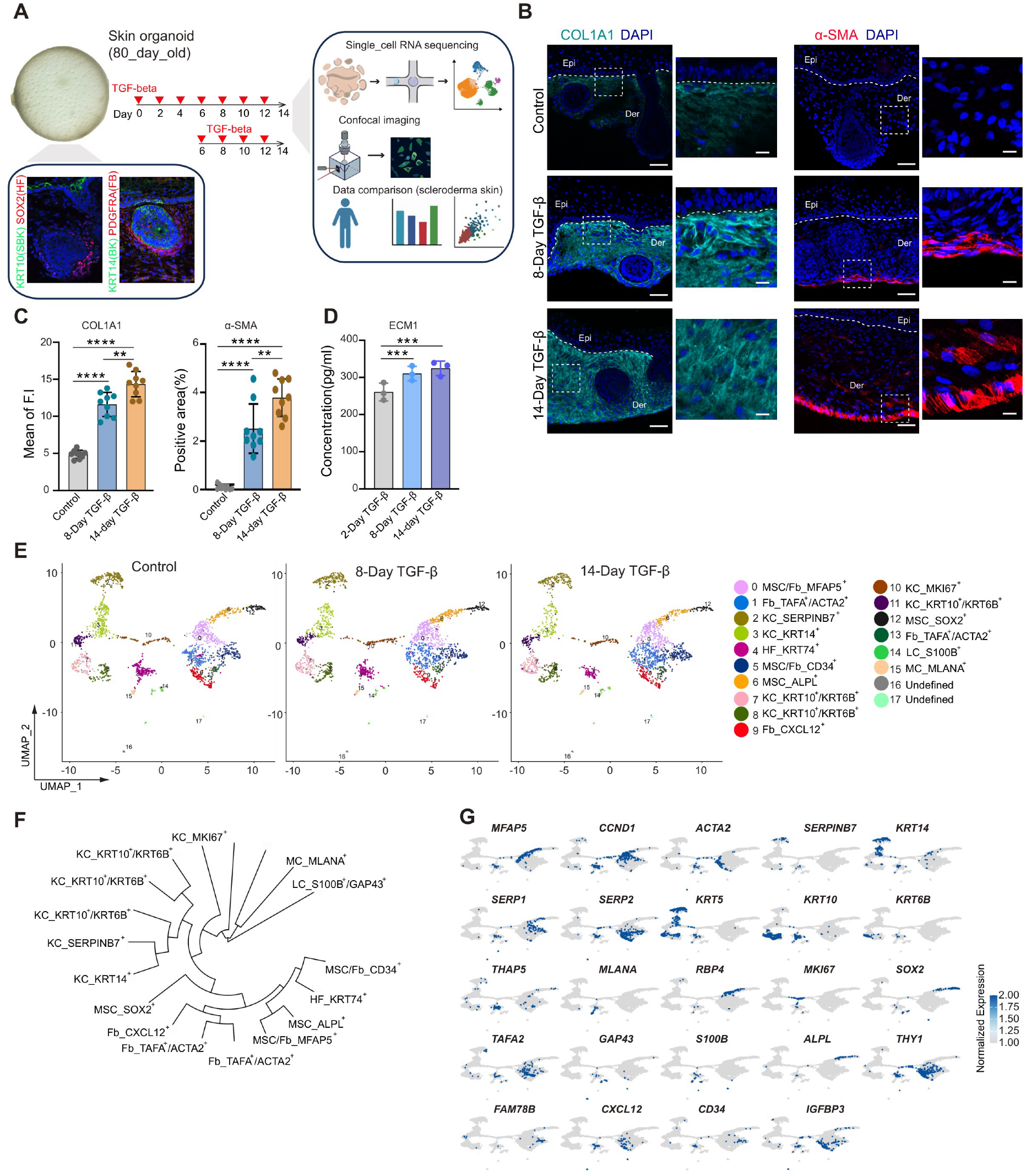
TGF-β induces fibrotic remodeling and cellular heterogeneity in human iPSC-derived skin organoids. (A) Schematic of the human iPSC-derived skin organoid system and experimental design. Skin organoids were generated and matured for 80 days before treatment. Cell identity and lineage specification were validated using canonical markers: KRT14 and KRT10 for epidermal keratinocytes, PDGFRA for dermal fibroblasts, and SOX2 for hair follicle–associated dermal condensate cells. TGF-β (10 ng/mL) was administered every other day for up to 14 days. To control for developmental stage–dependent effects, treatment was initiated at either day 0 or day 6, and all samples were collected at a common endpoint (day 14). Untreated organoids were included as controls. SBK: super basal keratinocyte; BK: basal keratinocyte; FB: fibroblast; HF: hair follicle. The graphical illustrations were created using FigureLab.ai. (B) and (C) Representative immunofluorescence images of COL1A1 and α-SMA in control and TGF-β–treated organoids following 8-day and 14-day treatment paradigms, with corresponding quantification of fluorescence intensity. For each condition, n = 3 independent organoids were analyzed, with 3 randomly selected regions per organoid. Epi, epidermis; Der, dermis. White dot line: boundary between epidermis and dermis. Scale bar, 100 μm. (D) Quantification of ECM1 protein levels in culture supernatants by ELISA at the indicated time points (2, 8, and 14 days) under control and TGF-β treatment conditions. n = 3 independent organoids per condition. (E) UMAP projection of integrated single-cell RNA-seq datasets from control and TGF-β–treated organoids (n = 8 pooled organoids per condition), revealing 17 transcriptionally distinct cell clusters. (F) Cyclic hierarchical clustering analysis depicting transcriptional relationships and continuity among the identified cell populations. (G) Feature plots showing the expression of representative marker genes used to annotate epidermal and dermal cell populations across clusters. Data in (C) and (D) are presented as mean ± SEM. One-way ANOVA determined statistical significance with multiple comparisons (\*\**P* < 0.01, \*\*\**P* < 0.001, \*\*\*\**P*<0.0001).

We found that TGF-β treatment induced progressive fibrotic remodeling marked by ECM accumulation and myofibroblast activation. Immunofluorescence (IF) analysis showed increased COL1A1 expression within the stromal compartment compared with untreated controls, with quantitative image analysis confirming significant elevation after 8 days and further increases by 14 days (Fig. 1B and C). Moreover, α-SMA staining showed an aligned organization in the deep dermal region after 8 days, accompanied by an increased number of α-SMA⁺ cells, and transitioned to a more intermingled pattern by 14 days, indicating expansion of myofibroblasts and progressive structural remodeling of the dermal compartment (Fig.1B and C). This transition is consistent with early wound-healing–like contractile organization, followed by disorganized matrix remodeling characteristic of fibrotic progression ^(27,28)^. These fibrotic changes were reproducible in an independent iPSC line, supporting robustness across genetic backgrounds (Supplementary Fig. 1A to B). To independently quantify ECM remodeling, we measured ECM1 levels in culture supernatants by ELISA and observed a significant increase over the course of treatment (Fig. 1D). Together, these data demonstrate that TGF-β induces fibroblast activation and ECM accumulation in iPSC-derived skin organoids.

To characterize cell-type composition and transcriptional changes during fibrotic remodeling, we performed single-cell RNA sequencing (scRNA-seq) on untreated control organoids and TGF-β–treated organoids collected after 8 or 14 days of stimulation. Following quality control and filtering, we obtained high-quality transcriptomic profiles from all three conditions for downstream analysis (Supplementary Fig. 1C). Integration and dimensionality reduction identified 17 transcriptionally distinct clusters spanning epidermal and dermal compartments, with all but clusters 16 and 17 annotated based on canonical lineage markers and cluster-enriched gene expression patterns (Fig. 1E). We resolved six fibroblast-related clusters, including a SOX2⁺ undifferentiated/early mesenchymal (MSC-SOX2^+^) population and multiple fibroblast states with distinct transcriptional programs. These included ALPL⁺ mesenchymal (MSC-ALPL^+^) cells, MFAP5⁺ mesenchymal/fibroblast-like (MSC/Fb-MFAP5^+^) cells, CD34⁺ mesenchymal/fibroblast-like (MSC/Fb-CD34^+^) cells, a TAFA2⁺/ACTA2 contractile fibroblast (Fb-TAFA^+^/ACTA2^+^) subset, and CXCL12⁺ signaling-associated fibroblasts (Fb-CXCL12^+^). In parallel, we identified keratinocyte populations spanning basal (KRT14⁺), transitional, and superbasal states (KRT10⁺ and KRT6B⁺), as well as MKI67⁺ proliferative and SERPINB7⁺ stress-associated keratinocytes, along with KRT74+ hair follicle lineage cells. Additional populations included S100B⁺ Langerhans cells and MLANA⁺ melanocytes, with feature gene expression supporting cluster annotation. We next performed cyclic hierarchical clustering, which revealed progressive transcriptional relatedness and state transitions across subpopulations (Fig. 1F). We then defined cluster-specific gene signatures across the integrated dataset (Supplementary Fig. 1D), using the top marker genes for each cluster (top 10 for heatmap visualization and top 50 for signature definition; Supplementary Table 1). Feature plots of representative markers further illustrated cell–type–specific expression patterns, supporting cluster annotation and highlighting distinct fibroblast and epithelial states (Fig. 1G). Collectively, these data delineate the transcriptional landscape of epidermal and dermal compartments and capture fibroblast heterogeneity during fibrotic remodeling in human iPSC-derived skin organoids.

### Dermal pseudotime reconstruction delineates fibroblast subpopulation state transitions and associated signaling program rewiring

To determine how TGF-β exposure altered cellular composition, we quantified cluster abundance in control and treated organoids. This revealed progressive remodeling across dermal and epidermal compartments (Fig. 2A). In the dermis, MSC-SOX2^+^ and MSC/Fb-MFAP5^+^ subpopulations remained largely unchanged, whereas fibrosis-associated fibroblast states expanded, including the Fb-TAFA2⁺/ACTA2⁺ contractile subset and MSC/Fb-CD34^+^ cells; notably, the Fb-CXCL12^+^ subpopulation was reduced, suggesting a shift toward a more contractile and ECM-remodeling fibroblast landscape. In the epidermis, basal, cycling, and hair follicle–associated keratinocytes decreased, whereas superbasal keratinocytes increased, consistent with altered differentiation dynamics and reduced regenerative capacity observed in fibrotic and pro-inflammatory skin remodeling contexts ^(29,30)^. Langerhans cells and melanocytes showed minimal changes, suggesting relative stability of these compartments, in line with reports that fibrosis-driven transcriptional alterations are predominantly fibroblast- and ECM-centered rather than broadly affecting epidermal immune or pigment cell lineages ^(31–34)^. We generated violin plots showing the expression of representative marker genes defining each fibroblast subcluster, including *SOX2*, *ALPL*, *MFAP5*, *CD34*, *CXCL12*, *ACTA2*, and *TAFA2*, together with classical fibroblast-associated genes such as *LAMC3*, *FN1*, and *SFRP1/2* across fibroblast subpopulations, highlighting the molecular heterogeneity and distinct transcriptional identities of each subset (Fig. 2B).

**Figure 2.**
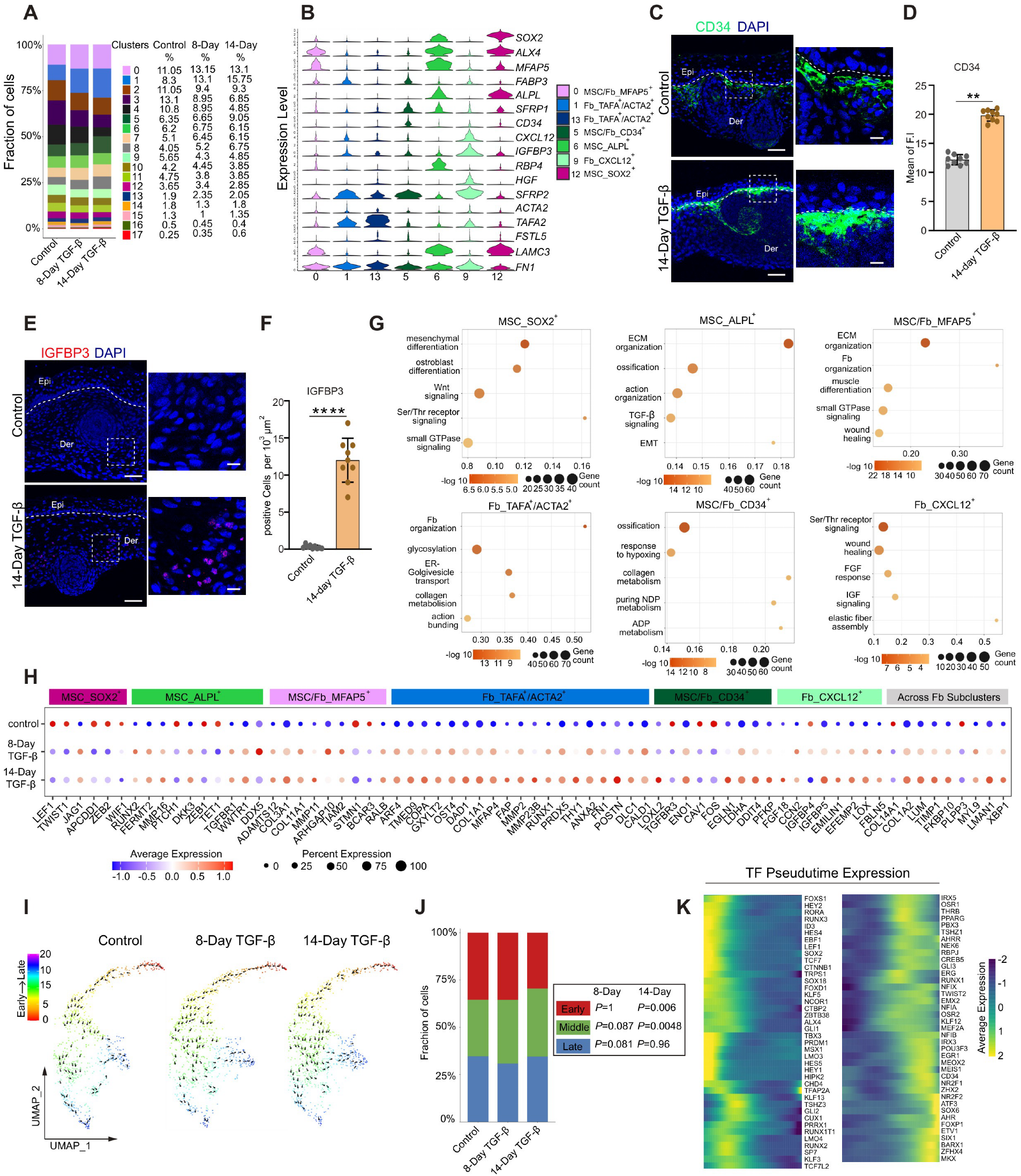
Fibrotic remodeling is accompanied by the emergence of transcriptionally diverse fibroblast subpopulations. (A) Relative abundance of cell clusters in control and TGF-β–treated organoids at 8-day and 14-day treatment time points. Proportional representation of major dermal and epidermal populations is shown for each condition, with quantitative comparison across time points. (B) Violin plots displaying expression levels of representative marker genes across annotated fibroblast subclusters. (C) to (F) IF staining of CD34 (C and D) and IGFBP3 (E and F) in control and TGF-β–treated organoids following 14-day treatment, with corresponding quantification of fluorescence intensity. For each condition, n = 3 independent organoids were analyzed, with 3 randomly selected regions per organoid. Epi, epidermis; Der, dermis. White dot line: boundary between epidermis and dermis. Scale bar, 100 μm. Scale bar, 100 μm. (G) Gene Ontology (GO) enrichment analysis of fibroblast subclusters (SOX2⁺, MFAP5⁺, CD34⁺, IGFBP3⁺, TAFA^+^/ACTA2⁺, and CXCL12^+^), identifying biological processes and functional programs associated with each cell state. (H) Bubble plot summarizing cluster-specific transcriptional responses to TGF-β treatment at 8-day and 14-day time points, with dot size representing the proportion of differentially expressed genes and color indicating relative expression changes across fibroblast populations. (I) Pseudotime vector field analysis illustrating directional transitions across fibroblast populations in control and TGF-β–treated organoids (8-day and 14-day conditions), capturing dynamic relationships among transcriptionally defined states. (J) Quantification of cell distribution across pseudotime-defined remodeling states, showing the relative proportion of cells assigned to each stage under control and treatment conditions. Statistical significance was assessed by Fisher’s exact test. (K) Heatmap of temporally ordered transcription factor (TF) modules across pseudotime, illustrating coordinated gene regulatory programs associated with early-, intermediate-, and late-stage transitions during TGF-β– induced remodeling. Data in (D) and (F) are presented as mean ± SEM. One-way ANOVA determined statistical significance with multiple comparisons (\*\**P* < 0.01, \*\*\*\**P* < 0.0001).

We further performed IF staining to validate marker expression and observed that CD34 expression significantly increased following 14-day TGF-β treatment (Fig. 2C and D; Supplementary Fig. 2A and B). In control organoids, MSC/Fb-CD34^+^ cells were primarily localized beneath the epidermal layer, whereas TGF-β treatment induced more intensive CD34 expression both within the subepidermal region and around hair follicle–like structures. These findings suggest that TGF-β signaling promotes spatial reorganization of CD34⁺ mesenchymal populations during fibrotic remodeling in this system. Given the developmental immaturity of skin organoids, these CD34⁺ populations likely represent heterogeneous mesenchymal compartments undergoing dynamic remodeling rather than fully mature adult dermal fibroblast subtypes, consistent with recent single-cell studies demonstrating functional heterogeneity of CD34⁺ stromal populations during skin homeostasis and wound repair ^(35,36)^.

Moreover, both scRNA-seq and qPCR demonstrated reduced *CXCL12* expression following TGF-β treatment in our skin organoid model (Supplementary Fig 2C and D). Because CXCL12 is associated with homeostatic fibroblast populations in the skin ^(37)^, its reduction likely reflects fibroblast state transitions during fibrotic remodeling. Interestingly, within the CXCL12⁺ fibroblast subcluster, *PDGFRL*, *CFH*, and *IGFBP3* expression increased following TGF-β treatment, whereas *HGF*, *EGFL6*, and *IGSF10* were progressively downregulated, indicating an activated stress-responsive phenotype (Supplementary Fig. 2D). Consistent with the scRNA-seq findings, IGFBP3 expression was markedly increased following TGF-β treatment at day 14 and was primarily localized to cells in the basal region of treated organoids, whereas little to no signal was detected in untreated controls (Fig. 2E and F; Supplementary Fig. 2E and F). These observations support the emergence of an IGFBP3-associated stromal population during fibrotic remodeling in this system ^(38)^.

We next performed Gene Ontology (GO) enrichment analysis, which revealed distinct functional programs across fibroblast subclusters (Fig. 2G). MSC-SOX2⁺ cells were enriched for mesenchymal differentiation and WNT signaling; MSC-ALPL^+^ population for extracellular matrix organization and TGF-β signaling; MSC/Fb-MFAP5⁺ cells for collagen organization and small GTPase signaling; TAFA^+^/ACTA2^+^ cells for Golgi transport and collagen metabolism; MSC/Fb-CD34^+^ subset for hypoxia response and metabolic processes; and Fb-CXCL12^+^ population for receptor kinase signaling and FGF pathways. These programs were supported by cluster-specific transcriptional changes relative to untreated controls, including reduced *LEF1* and *TWIST1* in SOX2^+^ cluster; increased *FERMT2*, *TGFBR1* and *MMP16* in ALPL^+^ cluster—genes associated with TGF signaling and matrix remodeling—alongside reduced *TET1* and *ZEB1*; upregulation of collagen remodeling genes (*ADAMTS12*, *COL3A1*, *COL11A1*, and *MMP11*) and altered small GTPase regulators (*ARHGAP10*, *TIAM2*, *STMN1*, *RALB*, and *BCAR3*) in MFAP5^+^ cluster; increased ER–Golgi and matrix-associated genes (*ARF4*, *TMED6*, *COPA*, *FAP*, *FN1*, and *POSTN*) in TAFA^+^/ACTA2^+^ cluster; hypoxia-related genes (*ENO1* and *CAV1*) in CD34+ cluster; and increased *FGF18* and *CCN2* with reduced *IGFBP4* and *BMP6/7* and upregulation of matrix-stabilizing genes (*EMILIN1*, *EFEMP2*, *LOX*, and *FBLN5*) in CXCL12^+^ cluster.

Signaling pathway analysis further revealed coordinated, state-specific remodeling across fibroblast subclusters in response to TGF-β (Supplementary Fig. 2G). EGFR signaling decreased in MSC/Fb-MFAP5⁺ cells but increased in the Fb-CXCL12⁺ cells, whereas JAK–STAT signaling was broadly elevated, and MAPK activity was generally reduced across multiple fibroblast states. PI3K signaling was decreased in the Fb-TAFA2⁺/ACTA2⁺ population, while WNT activity was enriched in MSC-SOX2⁺ cells. In contrast, NF-κB, TNF-α, and VEGF pathways were preferentially increased in MSC/Fb-CD34⁺ and Fb-CXCL12⁺ populations, consistent with inflammatory and angiogenic signaling programs. Together, these results highlight coordinated rewiring of signaling networks associated with fibroblast state diversification during fibrotic remodeling.

Pseudotime analysis revealed two bifurcating trajectories emerging from the MSC/Fb-MFAP5⁺ state (Supplementary Fig. 2H, Left panel). In Branch 1, MFAP5⁺ cells first transitioned toward the ACTA2⁺ contractile population and subsequently progressed to the CXCL12⁺ subset, indicating that the anti-fibrotic state arises downstream of contractile activation during fibroblast remodeling. In Branch 2, MFAP5⁺ cells progressed directly toward the CD34⁺ cluster. Notably, no distinct lineage bifurcation unique to either control or TGF-treated organoids was observed, suggesting preservation of overall fibroblast fate despite transcriptional remodeling (Supplementary Fig. 2H, right panel). To visualize directional dynamics, we constructed a pseudotime-based vector field on the UMAP embedding. Compared with control organoids, 14-day TGF-β–treated organoids exhibited a pronounced directional transition from ACTA2⁺ contractile cells toward CD34⁺ cells at a mid-pseudotime stage, accompanied by expansion of this population (*P*=0.0048, Fisher’s exact test) (Fig. 2I and J). Combined with the hypoxia-associated gene features observed in CD34⁺ cells (Fig. 3G), these findings suggest that contractile activation precedes engagement of hypoxia-related programs during fibrotic remodeling, linking myofibroblast activation to subsequent metabolic adaptation.

**Figure 3.**
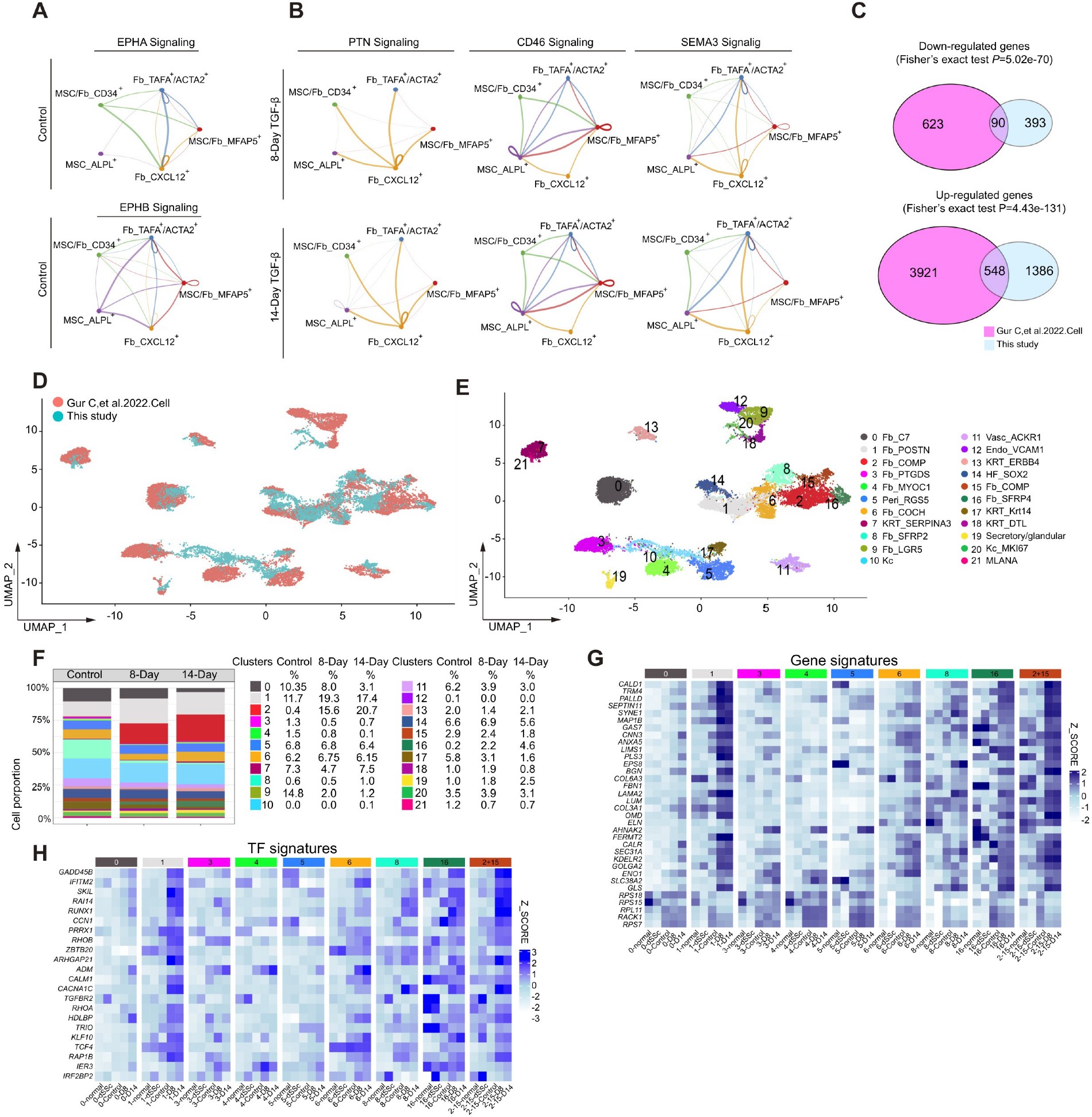
Fibrotic organoids exhibit extensive intercellular signaling rewiring and recapitulate the transcriptional programs observed in human skin scleroderma. (A, B) CellChat analysis of fibroblast–fibroblast communication networks in control and TGF-β–treated organoids under 8-day and 14-day treatment conditions. Network visualizations depict the relative contribution and distribution of top-ranked signaling pathways across fibroblast subclusters. (C) Comparative pseudo-bulk RNA analysis between fibrotic organoids and published single-cell RNA-seq datasets derived from human scleroderma skin. Overlap of differentially expressed genes was assessed to identify shared transcriptional programs associated with fibrotic remodeling across two systems. (D, E) Integrated UMAP visualization of Harmony-corrected datasets identified 22 shared transcriptionally distinct subclusters across skin organoids and human patient skin tissues (D). Cell type annotation, guided by the reference scleroderma dataset, identified major populations including fibroblast, epithelial, stromal vascular–associated, hair follicle– related, and melanocyte lineages, enabling cross-system comparison of cellular states (E). (F) Analysis of fibroblast subcluster composition across organoid datasets. (G, H) Heatmap showing comparative gene and TF signatures between scleroderma (normal and dSSc) and organoid (control, D8, and D14) datasets. Color intensity indicates relative expression levels across samples and conditions.

Furthermore, we revealed three temporally ordered transcription factor (TF) modules that evolved along with TGF-β treatment (Fig. 2K). Early pseudotime was enriched for progenitor regulators (*SOX2*, *LEF1*/*TCF7*, *CTNNB1*, *GLI1*, *FOXD1*, *ALX4*, and *TBX3*) and Notch factors (*HEY1*/*2*, and *HES4*/*5*). Mid-pseudotime was characterized by induction of osteogenic regulators (*RUNX2* and *SP7*) together with TFs associated with fibroblast specification and remodeling, including *OSR1*, *PRRX1*, *TWIST2*, *CREB5*, *RBPJ*, *GLI2/3*, *RUNX1*, *NFIX*, and *TCF7L2*. Late pseudotime was marked by TFs associated with terminal states, including regulators of mesenchymal identity (*NR2F2*, *SOX6*, *FOXP1*, *SIX1*, *BARX1,* and *MKX*), stress response (*ATF3* and *AHR*), and transcriptional regulation (*ETV1* and *ZFHX4*). These TF programs varied across fibroblast subclusters and, together with TGF-β treatment, defined fibrosis-associated transcriptional reprogramming. Representative TF expression patterns further support these stage-specific programs across fibroblast subclusters (Supplementary Fig. 2I).

### Cell-cell communication analysis and integration with human scleroderma datasets reveal conserved fibroblast states and signaling programs

To characterize changes in intercellular signaling during fibrotic remodeling, we performed CellChat analysis on single-cell transcriptomic data. EPHA/B signaling, prominent in control organoids, was no longer among the top-ranked pathways in fibrotic organoids, suggesting altered stromal coordination ^(39)^ (Fig. 3A). In contrast, fibrotic organoids exhibited three emergent top pathways at both day 8 and 14: PTN, CD46, and SEMA3. PTN signaling, linked to cell survival and angiogenic responses ^(40)^, was centered in the CXCL12⁺ cluster and extended across multiple fibroblast populations, indicating hub-like signaling behavior. CD46 signaling, associated with complement inhibition ^(41)^, and SEMA signaling, which mediates regulates neural guidance cues and cell–cell interactions ^(42)^, were prominently enriched fibroblast all subclusters, consistent with activation of immune- and intercellular communication-associated programs (Fig. 3B). ECM- and adhesion-related pathways, including COLLAGEN, FN1, and PERIOSTIN, were enriched in TGF-treated organoids, indicating enhanced matrix deposition and stromal remodeling (Supplementary Fig. 3A). Further analysis of epithelial–stromal communication revealed marked remodeling in fibrotic organoids. Contact-dependent pathway NECTIN was reduced between keratinocytes and fibrotic fibroblasts, indicating disruption of structured epithelial–dermal boundary signaling ^(43)^. In contrast, NOTCH signaling was increased and non-classic WNT signaling newly emerged following TGF-β treatment, consistent with reactivation of developmental and remodeling-associated communication programs between epithelial and stromal compartments (Supplementary Fig. 3B).

To evaluate disease relevance, we compared the organoid dataset with published single-cell datasets from human diffuse scleroderma skin ^(44)^. Comparative pseudo-bulk transcriptomic analysis demonstrated significant concordance between fibrotic organoids and scleroderma dermis (Fig. 3C, Supplementary Table 2). Among genes downregulated in fibrotic organoids, 90 of 483 (18.63%) overlapped with downregulated genes in scleroderma skin (Fisher’s exact test, *P* = 5.02e-70). Concordance was even more pronounced for induced genes: 548 of 1934 upregulated genes (28.33%) were shared between datasets (Fisher’s exact test, *P* = 4.43e-131). Together, these results indicate that the fibrotic organoid model robustly recapitulates scleroderma-associated transcriptional programs, with particularly strong preservation of upregulated pro-fibrotic signatures. Integration by using the Harmony pipeline resolved 22 shared subclusters across combined samples (Fig. 3D and Supplementary Fig. 3C and D). Based on annotations from the scleroderma reference dataset, we resolved nine fibroblast subclusters, together with seven epithelial populations, three stromal-associated vascular cell populations, hair follicle subpopulations, and melanocytes (Fig. 3E). Importantly, multiple fibroblast states were conserved between scleroderma dermis and fibrotic organoids, including c0-C7⁺ papillary fibroblasts, c1-POSTN⁺ and c2-COMP⁺ matrix-producing fibroblasts, c3-PTGDS⁺ inflammatory-associated fibroblasts, c4-MYOC1⁺ remodeling fibroblasts, c6-COCH⁺ structural fibroblasts, c8-SFRP2⁺ canonical dermal fibroblasts, and c16-SFRP4⁺ remodeling–associated fibroblasts. Compared to the reported human scleroderma datasets, we observed preserved remodeling trends, including expansion of c1-POSTN⁺ and c2-COMP⁺ fibroblasts, reduction of c4-MYOC1⁺ populations, and minimal changes in c6-COCH⁺ fibroblasts across fibrotic organoids (Fig. 3F) ^(44)^. In contrast, other fibroblast populations exhibited organoid-specific alterations, suggesting context-dependent remodeling under TGF-beta-driven conditions. Notably, c0-C7⁺ complement-associated and c3-PTGDS⁺ fibroblasts were reduced, whereas c16-SFRP4⁺ fibroblasts were expanded, indicating conserved pro-fibrotic remodeling and loss of homeostatic or immunoregulatory states.

Moreover, we observed shared transcriptional programs between the organoid and scleroderma datasets, marked by increased expression of genes involved in cytoskeletal contractility, ECM remodeling, ER–Golgi–mediated secretion, and metabolic stress responses, alongside reduced expression of ribosome biogenesis and translation-related genes, consistent with conserved fibroblast activation signatures in fibrosis (Supplementary Fig. 3E). Representative genes were mapped across fibroblast subclusters (Fig. 3G). *SYNE1*, *ANXA5*, *PLS3*, and *EPS8* were enriched in c1-POSTN⁺ and c6-COCH⁺ fibroblast clusters, reflecting cytoskeletal remodeling. Meanwhile, *COL6A3*, *LAMA2*, and *FERMT2* were elevated in c1-POSTN⁺ and c2-COMP⁺ subclusters, supporting enhanced ECM organization and focal adhesion signaling. Genes associated with ER–Golgi secretory activity (*CALR*, *SEC31A*, *GOLGA2*) were enriched in the c0-C7⁺ fibroblast population, whereas *ENO1* and *SLC38A2* were elevated in the c4-MYOC1⁺ subcluster, consistent with metabolic stress adaptation. In contrast, ribosome and translation-related genes were broadly downregulated in c1-POSTN⁺, c2-COMP⁺, c8-SFRP2⁺, and c16-SFRP4⁺ populations. We further identified conserved activation of co-regulatory networks in both datasets (Fig. 3H). Genes associated with TGF-β signaling and fibroblast activation, including *SKIL*, *TGFBR2*, *KLF10*, and *RUNX1*, were consistently upregulated in both datasets. Rho GTPase regulators and cytoskeletal components (*RHOA/B*, *ARHGAP21*, and *TRIO*), along with stress-responsive factors (*GADD45B*, *IER3*, *IFITM2*, and *IRF2BP2*), were similarly elevated. Mapping these regulators across fibroblast subclusters revealed enrichment of *RUNX1* and *ZBTB20* in c6-COCH⁺, c16-SFRP4⁺, and c2-COMP⁺ populations, whereas *TGFBR2* was broadly upregulated across fibroblast states. Together, these findings highlight conserved transcriptional and regulatory programs underlying fibroblast activation in the organoid model and human scleroderma.

### RUNX2 is involved in fibrotic remodeling in organoids and skin fibrotic diseases

Given the induction of the osteo-like regulator RUNX2 along the activated fibroblast trajectory during early mid-pseudotime, we next examined its role in fibrotic remodeling. We first assessed RUNX2 expression in published single-cell datasets from scleroderma and keloid tissues, where it was enriched in disease-associated fibroblast populations compared with normal skin ^(44–46)^ (Fig. 4A). Consistently, RUNX2 was upregulated in fibrotic organoids in our single-cell dataset, predominantly in clusters MFAP^+^ and TAFA^+^/ACTA2^+^ during treatment, and in cluster CXCL12^+^ at the 14-day time point. (Fig. 4B). IF staining further confirmed markedly increased RUNX2 expression in TGF-β-treated compared with untreated control across two independent iPSC-derived organoids (Fig. 4C and D; Supplementary Fig 4A and B). RUNX2 expression was also significantly increased following 48 h of TGF-β stimulation in both immortalized and primary human skin fibroblasts (hereafter referred to as Im-Fb and Pr-Fb, respectively). (Fig. 4E).

**Figure 4.**
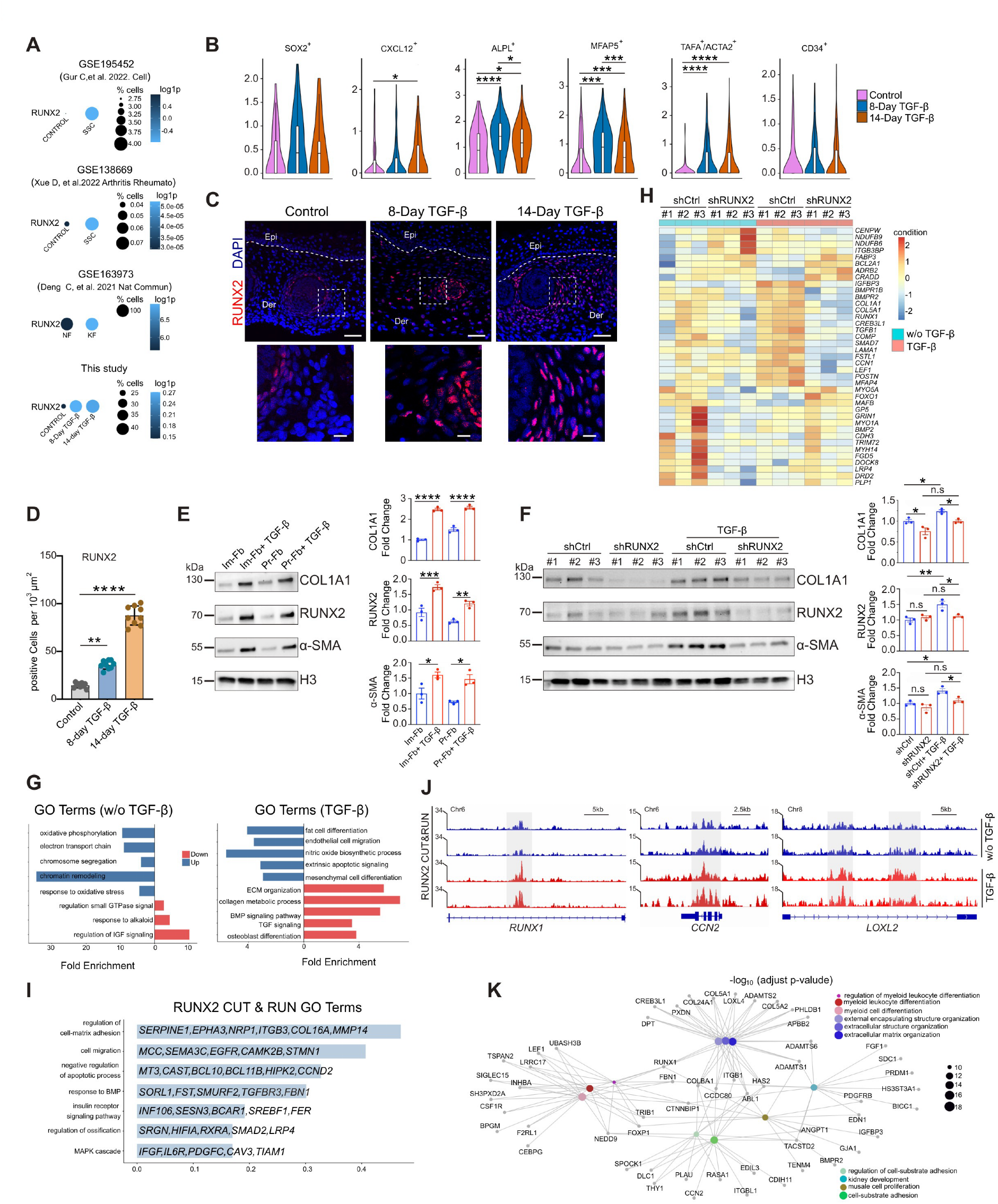
RUNX2 drives fibrotic remodeling and transcriptional regulation in TGF-β–treated skin fibroblasts. (A) Dot plot showing RUNX2 expression across fibroblasts in three published single-cell RNA sequencing datasets as indicated, alongside organoid-derived datasets. Dot size represents the proportion of RUNX2-expressing cells, and color intensity indicates relative expression levels. (B) Violin plot showing *RUNX2* expression across fibroblast subclusters from our scRNA dataset, illustrating the distribution and relative expression levels within each population. (C, D) IF staining of RUNX2 in control and TGF-β–treated organoids at 8-day and 14-day time points, with corresponding quantification of fluorescence intensity. For each condition, n = 3 independent organoids were analyzed, with 3 randomly selected regions per organoid. Epi, epidermis; Der, dermis. White dot line: boundary between epidermis and dermis. Scale bar, 100 μm. Scale bar, 100 μm. (E, F) RUNX2 expression in immortalized (Im-Fb) and primary (Pr-Fb) human skin fibroblasts (E), and validation of RUNX2 knockdown in three independent stable knockdown Im-Fb cell lines (F). Cells were serum-starved for 24 h and subsequently stimulated with TGF-β for 48 h. Protein expression and fibrosis-associated markers were assessed by Western blot. Quantification of protein expression levels was performed by densitometry and normalized to loading controls. (G) GO analysis of RUNX2 target genes under basal and TGF-treated conditions, showing enriched biological processes associated with RUNX2-mediated regulation. (H) Heatmap of differentially expressed genes following RUNX2 knockdown under TGF-β treatment conditions. (I) GO analysis of RUNX2-bound regions from CUT&RUN profiling highlights enrichment of biological processes and representative genes. (J) Representative IGV genome browser tracks showing RUNX2 occupancy at selected loci, including *RUNX1*, *CCN2*, and *LOXL2*. Highlighted regions (gray boxes) indicate loci with differential binding or regulatory changes across conditions. Data are representative of n = 2 biological replicates for each condition. (K) Gene–concept network analysis of RUNX2 target genes, highlighting interconnected functional modules. Data in panels (B), (D), (E), and (F) are presented as mean ± SEM. Statistical significance in panel (B) was determined using the Wilcoxon rank-sum test. Statistical significance in panels (D), (E), and (F) was determined using one-way ANOVA followed by multiple-comparison testing (\**P* < 0.05, \*\**P* < 0.01, \*\*\**P* < 0.001, \*\*\*\**P* < 0.0001).

To determine the functional role of RUNX2 in fibrotic remodeling, we generated stable RUNX2 knockdown (shRUNX2) and scrambled shRNA control Im-Fb cells and performed three independent biological replicates. Immunoblotting confirmed efficient RUNX2 depletion and demonstrated significantly reduced COL1A1 and α-SMA expression in TGF-β-stimulated cells across three independent shRUNX2 cell lines compared with scrambled shRNA controls (Fig. 4F). To define RUNX2-dependent transcriptional programs further, we performed bulk RNA sequencing under basal and TGF-β-stimulated conditions. Volcano plot analysis identified 145 upregulated and 353 downregulated genes in shRUNX2 cells under basal conditions (Benjamini–Hochberg-adjusted *P*<0.05,), whereas TGF-β treatment resulted in 412 upregulated and 233 downregulated genes relative to control cells (Benjamini–Hochberg-adjusted *P*<0.05) (Supplementary Fig. 4C and D; Supplementary Table 3 and 4). Under basal conditions, RUNX2 knockdown led to increased enrichment of pathways associated with the electron transport chain (*NDUFB6* and *NDUFB9*) and chromatin remodeling (*CENPW* and *ITGB3BP*). In contrast, pathways related to calcium metabolism (*DRD2*, *GP5*, *GRIN1*, and *PLP1*), actin organization (*MYH14*, *MYO1A*, and *MYO5A*), small GTPase signaling (*DOCK8*, *FGD5*, and *LRP4*), and insulin-like signaling (*BMP2*, *CDH3*, and *TRIM72*) were reduced. Strikingly, following TGF-β treatment, RUNX2 depletion promoted enrichment of fat cell lineage activity (*ADRB2*, *FABP3*, *FOXO1*, and *MAFB*) and apoptosis-related pathways (*BCL2A1*, *CRADD*) while key fibrotic programs—including ECM organization (*COL1A1*, *COL5A1*, *COMP*, *LAMA1*, *POSTN*, and *RUNX1*), collagen metabolic processes (*CREB3L1* and *MFAP4*), TGF-β/BMP signaling (*BMPR1B*, *BMPR2*, *FSTL1*, *SMAD7*, and *TGFB1*), and osteoblast differentiation (*CCN1*, *CREB3L1*, and *LEF1*)—were significantly suppressed (Fig 4 G and H).

RUNX2 is a well-established TF that regulates osteogenic differentiation ^(47)^. To investigate whether RUNX2 exerts a similar transcriptional regulatory role in activated fibroblasts and to identify its direct genomic targets, we performed CUT&RUN profiling in Pr-Fbs under basal and TGF-β–stimulated conditions. RUNX2 binding was enriched at promoter, gene body, and enhancer regions of genes involved in cell–matrix adhesion, ossification-related pathways, cell migration, anti-apoptosis, MAPK signaling, BMP response, and insulin receptor signaling (Fig. 4I; Supplementary Table 5). Together, these data support a direct role for RUNX2 in regulating transcriptional programs that drive fibroblast activation and fibrotic remodeling. Representative CUT&RUN profiles illustrate RUNX2 occupancy at key target loci, including *RUNX1*, a TF implicated in lineage specification and fibroblast plasticity; *CCN2*, a central mediator of ECM production and profibrotic signaling; and *LOXL2*, involved in ECM remodeling and tissue fibrosis (Fig. 4G). To identify direct RUNX2 target genes, CUT&RUN peaks from TGF-β–stimulated fibroblasts were intersected with genes downregulated in shRUNX2 cells following TGF-β treatment (bulk RNA-seq; FDR < 0.1 and log₂ fold change < 0). This integrative approach identified 233 direct RUNX2 targets, representing genes both bound by RUNX2 and transcriptionally dependent on its activity under fibrotic conditions (Supplementary Table 6). Gene–concept network analysis identified ECM organization, cell–substrate adhesion, and myeloid leukocyte differentiation as major RUNX2-regulated modules (Fig. 4K). *RUNX1*, *CCDC80*, *COL8A1*, *ABL2*, and *HAS2* emerged as central connector genes linking these pathways, suggesting that RUNX2 coordinates fibrotic remodeling through interconnected stromal and immune-associated regulatory networks.

### A Hybrid target–phenotype multi-task QSAR framework identifies candidate small-molecule inhibitors of RUNX2-driven fibrotic programs

Given the central role of RUNX2 in driving fibrotic remodeling, we prioritized RUNX2-dependent transcriptional programs for therapeutic intervention. To identify candidate inhibitors, we performed a pilot virtual screening of ∼113,000 commercially available compounds (Fig. 5A). Guided by available training data (e.g., >100 compounds per assay in the ChEMBL database), we selected 38 targets for modeling, including direct RUNX2-related targets identified in our study (e.g., BMPR1B, BMPR2, and LOXL2) (Supplementary Table 6) and additional fibrosis-associated targets (e.g., CK2, JAK3, TGFβ, HDAC9, HIF, EGFR, and VEGFR) ^(48–53)^. Quantitative structure-activity relationship (QSAR) models were constructed using multiple machine learning classifiers to predict inhibitory activity across these targets (see Methods for details). To capture compounds with potential indirect or multi-target antifibrotic effects, we further trained a phenotypic binary QSAR model on non-target-specific anti-fibrosis datasets using the same pipeline. From these analyses, 13 candidate compounds were selected to represent diverse predicted bioactivity profiles, all with acceptable drug-like properties and without known pan-assay interference structures (Fig. 5B; Supplementary Fig. 5A). We further screened those candidate compounds in TGF-β–stimulated human Pr-Fbs at 5 μM. Immunoblotting revealed that compound F0565-0303 significantly reduced RUNX2, COL1A1, and α-SMA expression, with no evidence of apoptosis, as indicated by the absence of cleaved caspase-3. (Fig. 5C).

**Figure 5.**
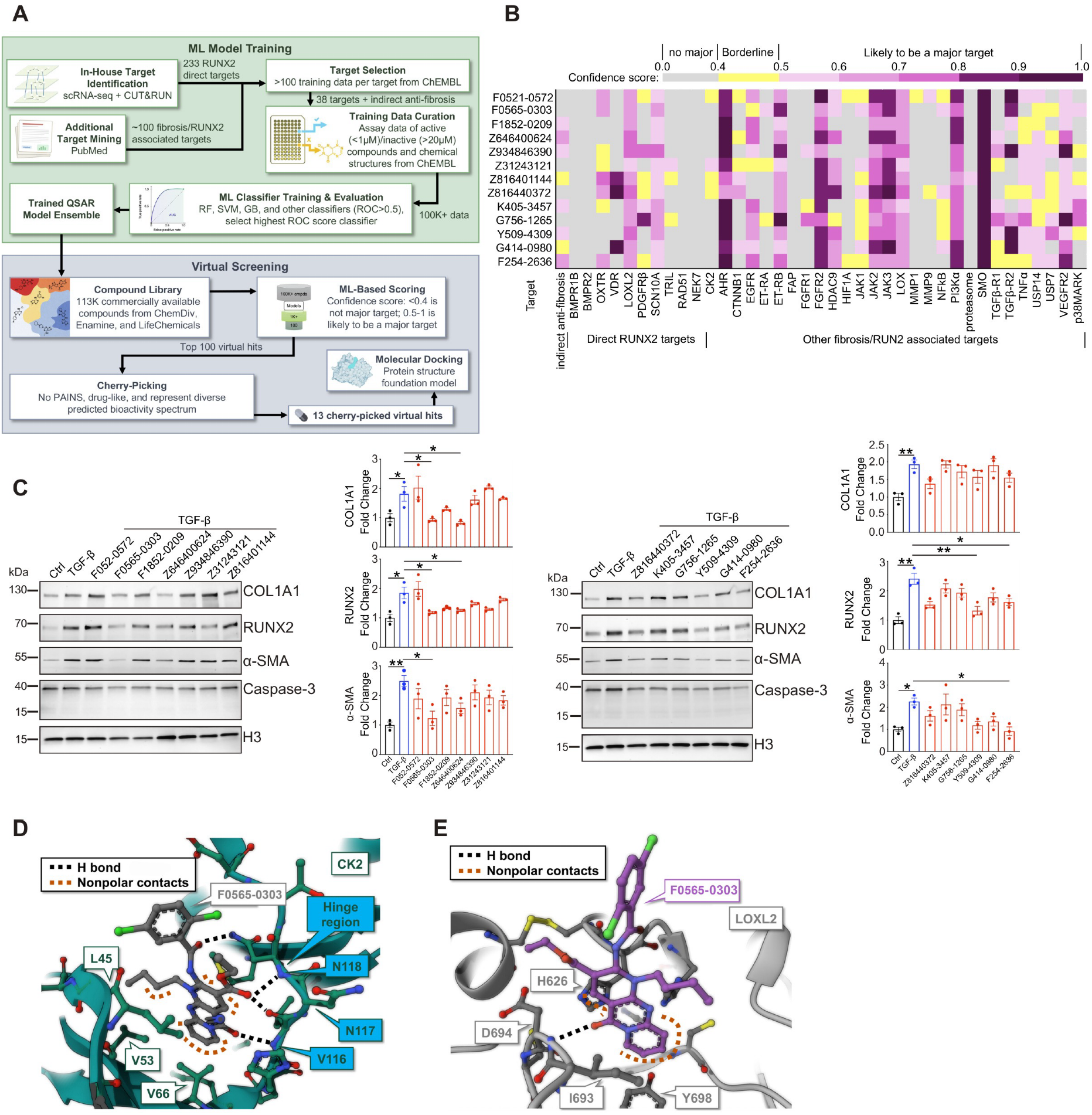
Polypharmacology-guided virtual screening identifies inhibitors of RUNX2 activity in fibrotic fibroblasts. (A) Schematic workflow of the polypharmacology-based machine learning model for compound prioritization, including model training, target prediction, and virtual screening of candidate molecules. (B) Virtual screening results for 13 selected candidate compounds. Predicted confidence scores are shown for each target–compound interaction, where a confidence score > 0.5 indicates a high likelihood of target inhibition. (C) Western blot analysis of RUNX2, COL1A1, and α-SMA in Pr-Fbs co-treated with TGF-β and candidate compounds (5 μM) for 48 h. Quantification of protein levels was performed by densitometry and normalized to loading controls. (D) Predicted binding mode of the compound F0565-0303 at human CK2α generated using a protein structure prediction framework. The compound is shown with gray carbon atoms, and kinase hinge region residues are highlighted in cyan. Predicted nonpolar interactions include hydrophobic contacts and π–π stacking. (E) Predicted binding mode of the compound F0565-0303 at human LOXL2. The compound is shown with lilac carbon atoms, with key interacting residues and predicted binding interactions indicated. Data in (C) are presented as mean ± SEM from three independent experiments. Statistical significance was determined by one-way ANOVA (\**P* < 0.05, \*\**P* < 0.01).

The CK2/USP7 pathway functions as a post-translational regulatory mechanism that stabilizes RUNX2 by preventing ubiquitin-dependent proteasomal degradation ^(47)^. Our machine learning model also identified USP7 as a potential target of F0565-0303 (Fig. 5B), prompting us to prioritize this compound for further functional studies. F0565-0303 exhibited an IC₅₀ of 9.1 μM and suppressed RUNX2 expression and fibrotic markers at 0.5 and 5 μM under TGF-β stimulation (Supplementary Fig. 5B and C). We further evaluated the effect of F0565-0303 on CK2α kinase activity. *In vitro* kinase assays showed that F0565-0303 reduced CK2α activity by approximately 20% at 20 μM relative to the other two candidate compounds, G756-1265 and Y509-4309 (Supplementary Fig. 5D), supporting a potential role for the CK2/USP7–RUNX2 axis in mediating its antifibrotic activity.

To explore potential modes of action of F0565-0303, we modeled putative binding interactions using the deep learning–based structural platform Boltz-2. Docking analysis predicted that F0565-0303 binds CK2α via hydrogen bonds with the hinge-region backbone and hydrophobic interactions (Fig. 5D), a binding mode characteristic of kinase inhibitors. We further investigated LOXL2 as an additional predicted target. LOXL2 was predicted to interact with F0565-0303 within its catalytic domain, including a π–π stacking interaction with residue Y689, suggesting potential inhibition of matrix stiffening (Fig. 5E). Overall, the mode of action of F0565-0303 was predicted to be binding to the hinge region of CK2α as a type-I kinase inhibitor with multiple other possible mechanisms such as binding to LOXL2 C-terminal to inhibit its catalytic function.

### Compound F0565-0303 suppresses RUNX2-associated fibrotic programs and attenuates dermal fibrosis *in vivo*

To further investigate the molecular basis of F0565-0303-mediated antifibrotic activity, we conducted transcriptomic profiling in human Pr-Fbs under four conditions: vehicle control, TGF-β, TGF-β plus F0565-0303, and F0565-0303 alone. Comparative analysis revealed that F0565-0303 treatment (TGF-β+F0565-0303 vs. TGF-β) partially reversed TGF-β–induced transcriptional changes (TGF-β vs. control; FDR < 0.05, |Log2FC| > 0.5). Specifically, F0565-0303 increased the expression of 92 genes that were reduced by TGF-β and decreased the expression of 65 genes induced by TGF-β (Supplementary Table 7). Genes increased by F0565-0303 were associated with oxidative stress response (*SOD2*), transcriptional regulation (*TCF7* and *ETS*), and chromatin remodeling (*HDAC9* and *NEAT1*), suggesting recovery of protective and homeostatic programs. In contrast, genes reduced by F0565-0303 were enriched for pathways related to ECM organization (*COL5A1*, *COL6A3*, *COL10A1*, *MFAP4*, and *SPARC*) and TGF-β signaling (*TGFBI* and *SEMA7A*), consistent with attenuation of fibrotic transcriptional programs (Fig. 6A; Supplementary Fig. 6A).

**Figure 6.**
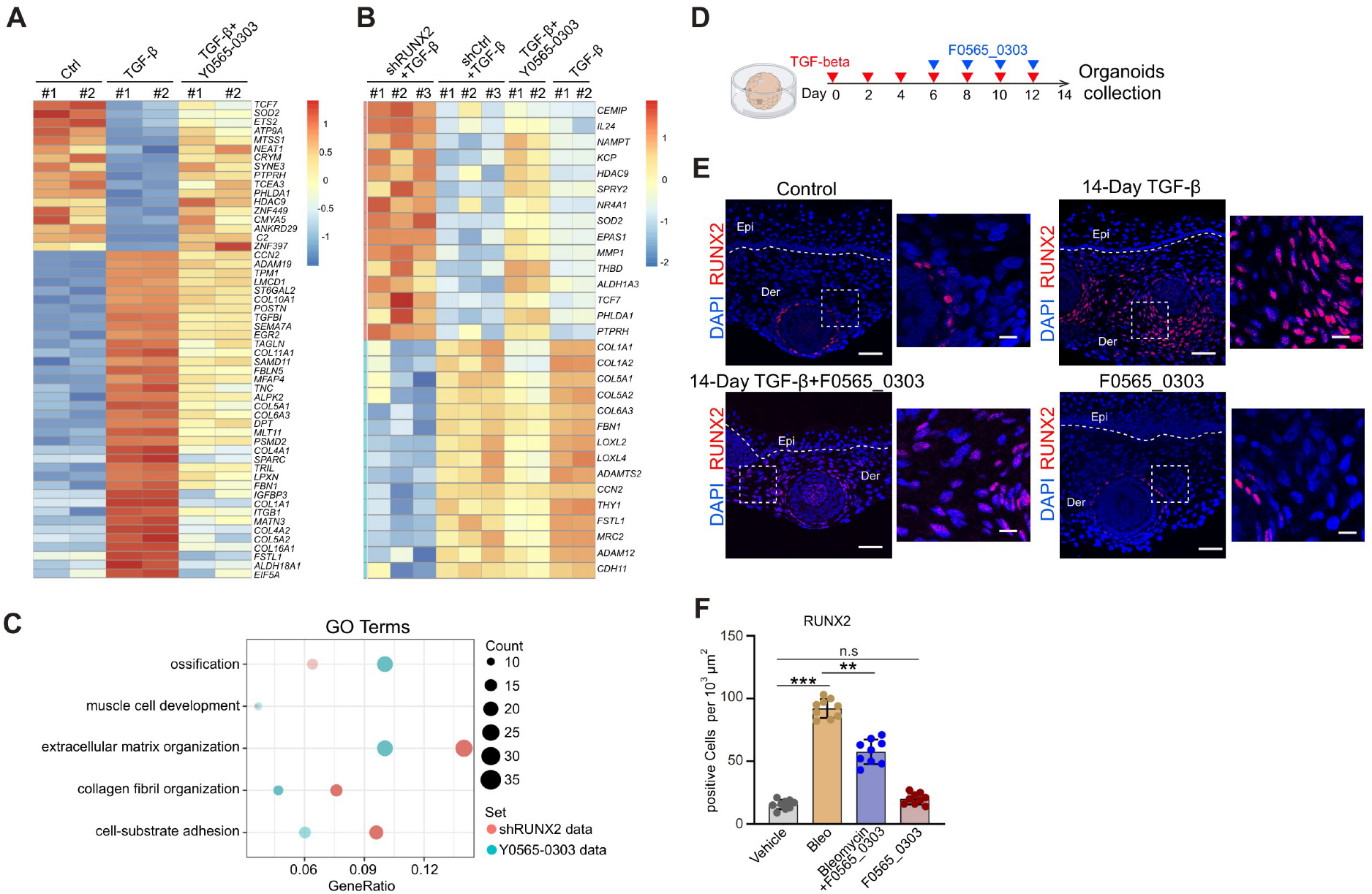
Pharmacological inhibition of RUNX2 by F0565-0303 suppresses fibrotic remodeling *in vitro*. (A) Heatmap of differentially expressed genes across experimental conditions, highlighting reversal of TGF-β–induced transcriptional changes by F0565-0303. (B) Heatmap of differentially expressed genes showing shared rescue of TGF-β– induced transcriptional changes by shRUNX2 and F0565-0303. (C) GO analysis for shared biological function between the knockdown of RUNX2 and F0565-0303 targets. (D) Diagram for Y0565-0303 treatment strategy using TGF-β-induced organoids. (E, F) IF staining of RUNX2 in human iPSC-derived skin organoids treated with TGF-β for 14 days, with 5 μM F0565-0303 co-treatment administered during the final 8 days of the treatment period. Quantification of fluorescence intensity is shown. For each condition, n = 3 independent organoids were analyzed, with 3 randomly selected regions per organoid. Epi, epidermis; Der, dermis. White dot line: boundary between epidermis and dermis. Scale bar, 100 μm. Scale bar, 100 μm. Data in (F) is presented as mean ± SEM. One-way ANOVA determined statistical significance with multiple comparisons (\*\**P* < 0.01, \*\*\**P* < 0.001).

To determine whether the transcriptional effects of F0565-0303 overlap with those induced by RUNX2 suppression, we compared RNA-seq profiles from shRUNX2 and F0565-0303-treated fibroblasts under TGF-β stimulation. This analysis identified shared reversal of TGF-β–induced transcriptional changes (FDR < 0.05, |log₂FC| > 0.5), including 32 genes increased, and 51 genes decreased in both conditions. Genes elevated in both datasets included *CEMIP*, *HDAC9*, *MMP1*, *TCF7*, and *PHLDA1*, which are associated with stress response and transcriptional regulation. In contrast, genes reduced in both conditions included key fibrotic and ECM components, such as *COL1A1*, *COL1A2*, *COL5A1*, *COL5A2*, *COL6A3*, *LOXL1/2*, *THY1*, *FSTL1*, and *ADAM12*, consistent with attenuation of profibrotic pathways (Fig. 6B and Supplementary Table 8). These overlapping genes were enriched for shared GO terms associated with ossification, ECM organization, and collagen fibril organization (Fig. 6C). Together, these findings indicate that F0565-0303 partially recapitulates the transcriptional effects of RUNX2 knockdown.

We next evaluated the effects of F0565-0303 in a skin organoid fibrosis model in which fibrotic organoids were first established by TGF-β treatment, followed by administration of compound F0565-0303 from day 8 to day 14 (Fig. 6D). IF staining demonstrated that F0565-0303 treatment significantly reduced RUNX2 expression in two independent iPSC-derived fibrotic organoid lines, further supporting suppression of RUNX2-associated fibrotic programs within a complex tissue environment (Fig. 6E and F; Supplementary Fig. 6B and C).

### Evaluation of F0565-0303 in the bleomycin-induced mouse model of skin fibrosis

Building on these findings, we next evaluated the *in vivo* antifibrotic efficacy of F0565-0303 using a bleomycin-induced mouse model of skin fibrosis. Mice were treated with bleomycin (3.0 mg/kg) for 14 days to induce dermal fibrosis, followed by administration of F0565-0303 (15 mg/kg) for an additional 14 days while fibrosis was maintained ^(54)^ (Fig. 7A). Histological analysis by H&E and Masson’s trichrome staining demonstrated that F0565-0303 treatment markedly attenuated dermal thickening and collagen accumulation compared with bleomycin-treated controls (Fig. 7B-E). In contrast, skin from mice treated with F0565-0303 alone exhibited histological features comparable to vehicle-treated controls, indicating no overt tissue abnormalities under basal conditions (Fig. 7B-E). IF staining further revealed significantly reduced RUNX2 and α-SMA expression in F0565-0303–treated mice relative to the bleomycin group, suggesting attenuation of profibrotic cellular response *in vivo* (Fig. 7F–I). Notably, the reduction in RUNX2 expression paralleled the attenuation of fibrotic features observed histologically, supporting a link between RUNX2 suppression and therapeutic response. Together, these results demonstrate that F0565-0303 effectively mitigates bleomycin-induced skin fibrosis and highlight RUNX2-dependent fibrotic programs as a promising therapeutic target for dermal fibrosis.

**Figure 7.**
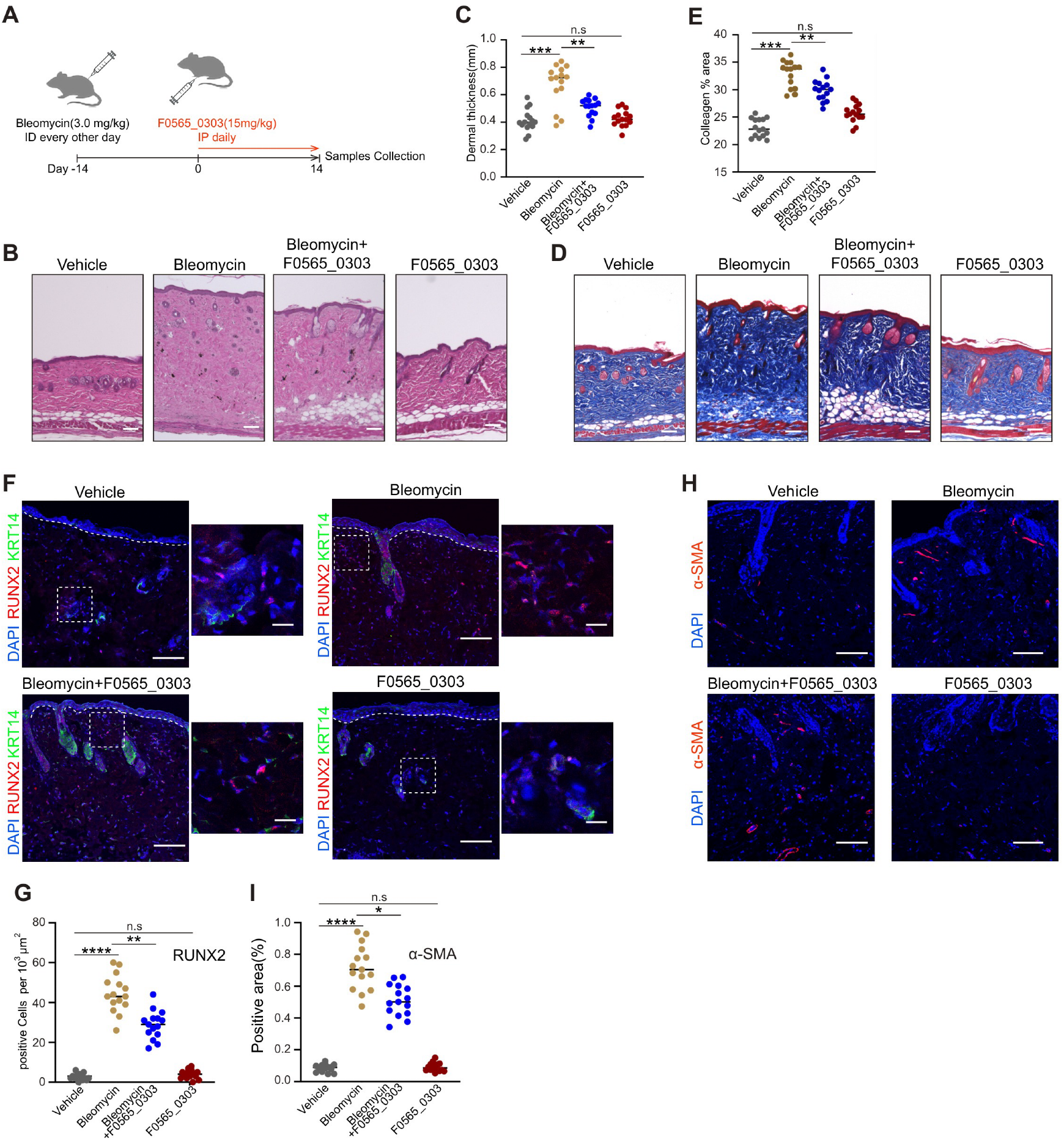
Therapeutic administration of F0565-0303 mitigates bleomycin-induced dermal fibrosis. (A) Schematic of the bleomycin-induced mouse skin fibrosis model and treatment timeline. (B to E) H&E staining (B, C) and Masson’s trichrome staining (D, E) of mouse skin sections from vehicle-, bleomycin-, bleomycin plus F0565-0303-, and F0565-0303 alone-treated mice showing dermal thickness and collagen deposition, with corresponding quantification. Scale bar, 200 μm. (F to I) IF staining of RUNX2 and α-SMA in mouse skin sections across the indicated conditions, with quantification of RUNX2- and α-SMA–positive cells. Scale bar, 200 μm. For all analyses, n = 5 mice per group, and 3 randomly selected regions were analyzed per mouse. Data in (C), (E), (G), and (I) are presented as mean ± SEM. One-way ANOVA determined statistical significance with multiple comparisons (\**P* < 0.05, \*\**P* < 0.01, \*\*\**P* < 0.001, \*\*\*\**P* < 0.0001).

## Discussion

We established a human iPSC-derived skin organoid model with TGF-β–induced fibrotic remodeling to investigate fibroblast state dynamics in a human-relevant system. Integrating single-cell transcriptomics with pseudotime trajectory analyses, we identified distinct fibroblast populations and demonstrated that fibrotic remodeling is driven by coordinated transitions across multiple functional states rather than uniform activation. Spatial analysis revealed dynamic organization of these states over time: at 8 days of treatment, α-SMA expression was largely restricted to basal regions, whereas by 14 days it expanded into a gradient extending toward apical compartments, indicating progressive propagation of myofibroblast activation. In contrast, CD34⁺ mesenchymal/fibroblast-like cells expanded following TGF-β treatment and became enriched in the apical region, defining spatially segregated fibroblast compartments along the fibrotic organoid axis. Notably, this spatial distribution differs from adult human skin, where CD34⁺ fibroblasts are typically localized within the reticular component ^(55)^. This discrepancy likely reflects the embryonic-like developmental origin and incomplete spatial maturation of organoid-derived dermal lineages, which may not fully recapitulate adult fibroblast lineage relationships and tissue architecture. Despite these architectural differences, integration with human scleroderma datasets revealed substantial conservation of fibroblast states and transcriptional programs, supporting the model’s relevance for studying core mechanisms of fibroblast activation and plasticity. Pseudotime and vector field analyses further supported dynamic interconversion between states rather than fixed lineage compartments. Together, these findings suggest that the organoid model recapitulates key features of fibrotic progression, including sustained myofibroblast activation, spatial reorganization of fibroblast populations, and fibroblast plasticity, while providing insight into conserved fibrotic programs operating within a developmentally derived human skin tissue context.

Beyond structural remodeling, we identified signaling-associated fibroblasts expressing IGF pathway components and chemokines, including IGFBP family members and CXCL12 (Fig. 2E, Supplementary Fig. 2C and D). Among these, IGFBP3 was specifically induced by TGF-β treatment, suggesting a fibrosis-associated signaling role within the IGF axis. Despite extensive transcriptional remodeling, this fibroblast population decreased in abundance following TGF-β treatment (Fig. 2E and F), consistent with functional reprogramming and potential transitions into other fibroblast states. Notably, *CXCL12* expression was significantly reduced during fibrotic remodeling, a finding consistent with reports that *Cxcl12* deletion promotes regenerative skin wound healing in postnatal mice ^(56)^, suggesting that CXCL12 may help regulate the balance between fibrosis and regeneration.

Our data also reveal coordinated reprogramming of stromal immune-associated pathways during fibrosis. C7 fibroblasts were reduced in TGF-β–treated organoids, whereas CD46 signaling was increased across fibrotic fibroblast states (Fig. 3B). Given that CD46 functions as a complement-inhibitory receptor ^(41)^, this inverse relationship suggests a shift from complement-associated fibroblast states toward complement-inhibitory programs during fibrotic remodeling. Additionally, PTGDS⁺ fibroblasts were reduced in our model, in contrast to prior reports describing increased PTGDS expression in scleroderma ^(44)^ (Fig. 3F). This discrepancy may reflect context-dependent fibroblast state dynamics during fibrotic remodeling. In our organoid system, the reduction of PTGDS⁺ fibroblasts following TGF-β treatment suggests selective remodeling of inflammatory- or immune-associated stromal populations during the transition toward matrix-producing profibrotic states. Thus, PTGDS⁺ fibroblasts may represent a homeostatic or immune-regulatory population that is depleted or reprogrammed during fibrosis progression. Alternatively, differences between organoid and patient tissue contexts, including the absence of immune and vascular components, may contribute to altered PTGDS-associated signaling and fibroblast composition.

Interestingly, under basal conditions, RUNX2 expression in organoids was relatively restricted and primarily localized around hair follicle–associated regions within the dermal compartment, consistent with previous studies implicating RUNX2 in hair follicle development, particularly within dermal papilla and perifollicular dermal sheath compartments involved in epithelial–mesenchymal signaling and follicular homeostasis ^(57)^. In contrast, TGF-β treatment induced broad expansion of RUNX2 expression throughout the dermis, accompanied by increased fibroblast activation and ECM– remodeling programs identified by single-cell analysis, suggesting that fibrotic signaling reprograms RUNX2 activity from localized homeostatic niches toward widespread stromal activation during fibrosis.

We identify RUNX2 as a regulator of fibroblast activation and fibrotic remodeling, extending its established function in osteogenic lineage specification to broader mesenchymal reprogramming within the fibrotic dermal microenvironment. The widespread induction of RUNX2 during TGF-β–driven fibrosis, together with suppression of fibrotic markers following RUNX2 depletion, supports its involvement in coordinating fibroblast activation and extracellular matrix remodeling programs (Fig. 4G and H). CUT&RUN profiling further delineated RUNX2 genomic target modules associated with extracellular matrix organization, cell–matrix adhesion, migration, and fibroblast activation (Fig. 4I). These analyses identified RUNX2 occupancy at multiple fibrosis-associated loci, supporting a direct contribution of RUNX2 to pathogenic stromal transcriptional programs. Notably, RUNX2 occupancy at the *RUNX1* locus suggests that RUNX2 acts upstream of RUNX1 to establish a profibrotic transcriptional program. Consistent with this model, *RUNX2* expression was enriched in the MSC/Fb-MFAP5⁺ population, whereas *RUNX1* was predominantly detected in the downstream Fb-TAFA2⁺/ACTA2⁺ contractile, MSC/Fb-CD34⁺, and Fb-CXCL12⁺ fibroblast states (Supplementary Fig. 2I), supporting a hierarchical RUNX2–RUNX1 regulatory axis during fibroblast activation.

Recent single-cell epigenomic studies in diffuse cutaneous systemic sclerosis have nominated RUNX2 as a putative regulatory transcription factor associated with pathogenic fibroblast populations based on chromatin accessibility analyses ^(44)^. Consistent with a profibrotic role, RUNX2 has been shown to regulate collagen gene expression in keloid fibroblasts ^(58)^. Beyond skin fibrosis, RUNX2 has also been implicated in pulmonary fibrosis, where it promotes pathological alveolar transitional cell states that contribute to disease progression ^(59)^.

Current FDA-approved therapies for systemic sclerosis, including nintedanib and tocilizumab ^(60,61)^, primarily target systemic sclerosis–associated interstitial lung disease, while no effective treatment for cutaneous fibrosis exists. Cutaneous fibrosis is largely managed by immunosuppressive agents that incompletely reverse established fibrotic remodeling. Consequently, finding a drug that can block skin fibrotic pathogenesis by directly targeting fibroblast activation and ECM deposition will provide a significant advance in this skin disease management. In this study, our discovery that a small molecule, F0565-0303, suppresses RUNX2-associated profibrotic programs and attenuates fibrosis in human skin fibroblasts as well as in a bleomycin-induced mouse model of fibrosis indicates a likely breakthrough and a step closer to clinical translation. Unlike previous approaches that directly inhibit RUNX2 transcriptional activity, such as the RUNX2 inhibitor CADD522 ^(62)^, our findings support modulation of RUNX2-associated remodeling networks as an important therapeutic strategy to suppress fibroblast activation and profibrotic signaling. It is known that direct targeting of transcription factors for therapeutic benefits often fails. Together, these results establish RUNX2 as a potentially actionable regulator of fibrotic remodeling and support further development of F0565-0303 for the treatment of cutaneous fibrosis in systemic sclerosis.

In summary, our study establishes a human iPSC-derived skin organoid model of fibrosis and identifies RUNX2 as a central regulator of fibroblast activation and ECM remodeling during TGF-β–driven fibrosis. Integrated single-cell, functional, and pharmacologic analyses reveal dynamic stromal reprogramming and induction of RUNX2-associated transcriptional programs during fibrotic progression. Together, these results highlight the utility of human skin organoid models for dissecting stromal regulatory networks and provide a framework for developing targeted interventions for fibrotic skin disease.

### Limitations and future directions

Several limitations should be considered. First, the organoid system lacks immune and vascular components that contribute to fibrosis *in vivo*, which may influence fibroblast– microenvironment interactions. Second, while CUT&RUN identifies RUNX2 genomic binding sites, further integration with functional assays will be necessary to define causal regulatory relationships and downstream transcriptional networks. Finally, although AI-mediated screening provides a powerful approach for candidate discovery, additional *in vivo* validation and optimization will be required to establish therapeutic efficacy and specificity.

## Acknowledgements

We thank the Sequencing Core Facility at UAB for their technical assistance.

## Author contributions

Conception, design, and supervision, S.Z., M.A., and L.J.; acquisition and analyses, J.K., S.M., F.A.A., M.W., S.M., S.M., S.L., M.W., J.A., Z.L., D.C.; Interpretation of data, F.A.A., S.M., S.Z., L.J.; experimental advice: M.K., J.C., C.R., C. E.; writing-original draft, S.M., S.Z., L.J.; writing-review & editing: S.Z., M.A., and L.J.

## Declaration of no conflict of interest

The authors declare that they have no competing interests.

## Data and materials availability

The processed data of this study have been deposited in the GEO database under accession codes GSE330906 (CUT&RUN datasets), GSE330910 (bulk RNA-seq datasets), and GSE330913 (scRNA-seq datasets). QSAR models for virtual screenings can be accessed via https://github.com/SZ-Lab7/Hybrid-target-phenotype-virtual-screening-RUNX2-pathway-fibrosis-.git. All data, codes, and materials used in the analysis are available within the Article, Supplementary Materials, or from the corresponding author upon reasonable request.

## Methods

### Cell lines and culturing conditions

Human induced pluripotent stem cell (hiPSC) lines, hereafter referred to as ATCC-hiPSC and STEMCELL-hiPSC, were purchased from ATCC (Cat #ACS-1027) and STEMCELL Technologies (Cat #200-0510), respectively, and cultured on Vitronectin (Gibco, Cat #A14700)-coated 6-well tissue culture plates. ATCC-hiPSC was used for all key experiments, and STEMCELL-hiPSC for validation. Cells were passaged every 3–4 days using Accutase (Gibco, #A11100501) according to the manufacturer’s instructions. Cultures were maintained in a humidified incubator at 37°C with 5% CO₂. Immortalized human skin fibroblasts were purchased from ABM (Cat #T0302), and primary adult human dermal fibroblasts were obtained from ATCC (Cat # PCS-201-012). Fibroblasts were cultured in Dulbecco’s Modified Eagle Medium (Hyclone #SH30243.01) supplemented with 10% fetal bovine serum (FBS, R&D Systems, Cat #11150) and 1% penicillin-streptomycin in a humidified incubator at 37°C with 5% CO₂.

### Animal care and use

Male C57BL/6J mice (10–12 weeks of age) were purchased from The Jackson Laboratory (Cat #000664) and housed under specific pathogen-free conditions with ad libitum access to food and water. Animals were maintained on a 12-hour light/dark cycle in a temperature- and humidity-controlled facility. All animal procedures were approved by the Institutional Animal Care and Use Committee (IACUC) at the University of Alabama at Birmingham (Protocol No. IACUC-23242) and were conducted in accordance with institutional guidelines for the care and use of laboratory animals.

## METHOD DETAILS

### Differentiation of hiPSC to skin organoids and treatment with TGF-β

Human skin organoids were generated from hiPSCs as previously described with minor modifications (17). Briefly, hiPSCs were dissociated into single cells using Accutase (Gibco, Cat# A1110501) and aggregated in ultra-low attachment 96-well plates on day 0 in Essential 6 medium (Gibco, Cat# A1516401) supplemented with Normocin (InvivoGen, Cat# ANT-NR-1) and embedded in 2% (v/v) growth factor–reduced Matrigel (Corning, Cat# 354230). Surface ectoderm induction was initiated by supplementation with SB431542 (10 μM, Stemgent, Cat# 04-0010-05), BMP4 (10 ng/mL, R&D Systems, Cat# 314-BP-010/CF), and bFGF (4 ng/mL, PeproTech, Cat# 100-18B). On day 3, patterning was further promoted in Essential 6 medium supplemented with bFGF (250 ng/mL) and LDN-193189 (1 μM, Stemgent, Cat# 04-0074-02). From day 12, organoids were transferred to floating culture in maturation medium consisting of a 1:1 mixture of Advanced DMEM/F12 (Gibco, Cat #12634010) and Neurobasal medium (Gibco, Cat# 21103049) supplemented with GlutaMAX (Gibco, Cat# 35050061), N2 (Gibco, Cat# 17502048), B27 (Gibco, Cat# 17504044), 2-mercaptoethanol (Gibco, #21985023), and Normocin, while maintaining 1% (v/v) Matrigel support as needed. From day 18 onward, organoids were cultured in maturation medium without Matrigel, with medium changes every 2–3 days. During long-term culture, organoids underwent self-organization and developed stratified epidermal and dermal-like compartments characteristic of developing human skin and were maintained for up to 80 days. Fibrotic changes were induced by treating skin organoids with TGF-β (10 ng/mL; Sigma-Aldrich, Cat# T7039) in maturation medium every other day for up to 14 days, with untreated organoids serving as controls. To account for potential developmental stage–dependent effects, TGF-β treatment was initiated at either day 0 (14-day exposure) or day 6 (8-day exposure). All organoids were collected at a unified endpoint on day 14 for downstream analyses, embedded in OCT compound, cryosectioned, and stored at −80°C until further use.

### ECM1 ELISA assay

Human ECM1 protein levels were quantified using the Human ECM1 SimpleStep ELISA Kit (Abcam, Cat# ab246524) according to the manufacturer’s instructions. Briefly, conditioned media were collected from organoid cultures at day 2, day 8, and day 14. Standards and samples were added to a 96-well plate pre-coated with capture antibody, followed by the addition of an antibody cocktail containing the detection antibody. After incubation at room temperature for 1 hour, wells were washed, and TMB substrate was added. The reaction was stopped using the provided stop solution, and absorbance was measured at 450 nm using a Synergy Neo2 multi-mode plate reader (BioTek). ECM1 concentrations were calculated from a standard curve generated using the supplied standards, with samples diluted as necessary to ensure values fell within the linear range of the assay. Data acquisition was performed using Gen5 software (version 3.00).

### Quantitative RT-PCR (qRT-PCR)

Total RNA was extracted using TRIzol reagent (Invitrogen, Cat# 15596026) according to the manufacturer’s instructions. Complementary DNA (cDNA) was synthesized using the SuperScript III First-Strand Synthesis System (Invitrogen, Cat#18080051) with oligo(dT)20 primers. Quantitative PCR (qPCR) was performed using TaqMan™ Fast Advanced Master Mix (Applied Biosystems, Cat# 4444557) on a QuantStudio 12K Flex Real-Time PCR System (Applied Biosystems). The thermal cycling conditions were as follows: initial denaturation at 95°C for 20 s, followed by 40 cycles of 95°C for 3 s and 60°C for 30 s.

### Establishment of stable RUNX2 knockdown fibroblasts and TGF-β treatment

Stable RUNX2 knockdown fibroblast lines were generated using human immortalized dermal fibroblasts transduced with lentiviral shRNA targeting RUNX2 (Santa Cruz Biotechnology, Cat# sc-37145-V). A non-targeting control shRNA lentivirus (Santa Cruz Biotechnology, Cat# sc-108080) was used as a control. Cells were cultured in DMEM supplemented with 10% fetal bovine serum and 1% penicillin-streptomycin throughout the procedure. Following transduction, cells were selected with puromycin (2 μg/mL, Gibco, Cat# A1113803) to establish three independent stable RUNX2 knockdown populations.

### Immunofluorescence (IF) staining

For IF staining of paraffin-embedded tissue sections, slides were deparaffinized in xylene and rehydrated through a graded ethanol series (100%, 95%, and 70% ethanol), followed by rinsing in deionized water. Antigen retrieval was performed using antigen unmasking solution (Vector Laboratories, Cat# H-3300) with microwave heating, followed by cooling to room temperature. Sections were then blocked with 5% goat serum in PBS for 45 min at room temperature, and primary antibodies diluted in 1% BSA in PBS were applied overnight at 4°C. For IF staining of cryosections, sections were blocked with 5% goat serum in TPBS (PBS containing 0.1% Triton X-100) for 45 min at room temperature, and primary antibodies diluted in 3% goat serum in TPBS were applied overnight at 4°C. The following day, sections were washed three times with PBS (10 min each), incubated with appropriate fluorophore-conjugated secondary antibodies for 1 h at room temperature, and washed again. Nuclei were counterstained with VECTASHIELD Antifade Mounting Medium with DAPI (Vector Laboratories, Cat# H-1200), and coverslips were mounted for imaging. Images were acquired using an FV3000 confocal laser scanning microscope (Olympus, USA) equipped with an FV3000 Galvo scan unit. Image acquisition was performed using FV31S-SW software (version 2.3.2.169), and identical imaging settings were applied across all experimental groups for quantitative comparisons. Primary and secondary antibodies are listed in the Key Resources Table.

### Fibroblast culture and treatment conditions

Skin fibroblasts were seeded in 6-well plates at a density of 0.8 × 10⁵ cells per well and cultured in Dulbecco’s Modified Eagle Medium (DMEM) supplemented with 10% fetal bovine serum (FBS) for 24 h. Cells were subsequently serum-starved in serum-free DMEM for an additional 24 h before treatment. For TGF-β stimulation experiments, cells were treated with 10 ng/mL TGF-β1 for 48 h. For compound treatment experiments, hit compounds were dissolved in dimethyl sulfoxide (DMSO) as 10 mM stock solutions and diluted in culture medium to a final concentration of 5 μM. Cells were co-treated with hit compounds and 10 ng/mL TGF-β1 for 48 h. Vehicle control cells received an equivalent volume of DMSO. Following treatment, cells were harvested for downstream analyses. Compounds are listed in the key resources table.

### Western-blot assays

Following treatment with TGF-β and/or small molecules at the indicated concentrations for 48 h, fibroblasts were washed three times with PBS and harvested in ice-cold PBS. Cells were pelleted by centrifugation at 300–500 × g for 5 min at 4°C. Cell pellets were lysed in ice-cold RIPA buffer (Santa Cruz Biotechnology, Cat #sc-249489) supplemented with protease and phosphatase inhibitor cocktails according to the manufacturer’s instructions. Lysates were incubated on ice for 30 min with intermittent vortexing to ensure complete lysis and homogenization. Samples were then centrifuged at 12,000 × g for 15 min at 4°C, and the supernatants were collected. Protein concentrations were determined using the Bio-Rad DC Protein Assay kit (Bio-Rad Laboratories, Cat #5000112). Equal amounts of protein (20 μg per sample) were mixed with 4× reducing Laemmli sample buffer and denatured at 95°C for 5 min. Proteins were separated by SDS-PAGE and transferred onto polyvinylidene difluoride (PVDF) membranes. Membranes were blocked in 5% nonfat dry milk prepared in Tris-buffered saline containing 0.1% Tween-20 (TBS-T) for 1 h at room temperature. Following blocking, membranes were incubated overnight at 4°C with primary antibodies diluted in 5% nonfat dry milk at the manufacturer’s recommended concentrations. The following day, membranes were washed three times with TBS-T for 10 min each and incubated with the appropriate horseradish peroxidase-conjugated secondary antibodies diluted in 5% nonfat dry milk for 2 h at room temperature with gentle rocking. Membranes were subsequently washed three times with TBS-T for 10 min each. Protein bands were visualized using enhanced chemiluminescence (ECL) substrate (Thermo Fisher Scientific, Cat #32106) and imaged using an Invitrogen iBright FL1500 Imaging System (Thermo Fisher Scientific). Band intensities were quantified using ImageJ software (National Institutes of Health, Bethesda, MD, USA) and normalized to the corresponding endogenous loading controls. Details of all primary and secondary antibodies used in this study are provided in the Key Resources Table.

### Machine learning-based Virtual Screening and binding mode prediction

The 233 RUNX2 direct targets in this paper and ∼100 known RUNX2- or fibrosis-associated targets from the literature were evaluated for the feasibility of developing ligand-based QSAR models. Targets that had >100 compounds with assay data as training data were processed for model training. Data were collected from the ChEMBL database. For a given target, multiple categorical models were trained using various machine learning classifiers, including SVM, random forests, and gradient boosting. IC50 <1 μM and >20 μM were used to define active and inactive samples, respectively. The model with the highest ROC score was retained as the final QSAR model for that target. Only models with ROC scores >0.85 were retained. The phenotypic fibrosis QSAR model was built with the same approach and training data as the anti-fibrosis assay data of 1,102 compounds. Eventually, QSAR models were built for 38 targets plus the anti-fibrosis phenotype. A total of 113K commercially available compounds were collected from catalogs of vendors ChemDiv, Enamine, and LifeChemicals. The SMILES strings representing the chemical structures of those compounds were used as input for virtual screening across all 39 QSAR models. Compounds predicted as positive with a confidence score >0.5 were considered to have a likelihood of inhibitory activity. The training data, training results, final model versions, and the virtual screening compound library are available in the GitHub repository (https://github.com/SZ-Lab7/Hybrid-target-phenotype-virtual-screening-RUNX2-pathway-fibrosis-.git). Virtual hits were checked against the Pan-Assay-Interference Structure (PAINS) filter. None of the virtual hits contained PAINS. To aid cherry-picking, the drug-likeness of virtual hits was evaluated via a combination of drug-likeness scores ^(75)^ and visual inspection of the chemical structures. Target binding poses of the virtual hits were generated using the protein structure foundation model Boltz-2. The SMILES of the compounds and the sequences of the corresponding human protein targets from the UniProt database were used as the input for Boltz-2 ^(76)^. Multi-sequence alignment was used to improve the accuracy of protein conformation. Physics-based potentials were used to steer predictions at inference time to improve the physicality of the generated poses. Binding poses were visualized with the open-source molecular visualizer Molstar (https://molstar.org).

### Cell Viability Assay

Cell viability was determined using the MTT Cell Proliferation Assay Kit (Abcam, Cat# ab211091) according to the manufacturer’s instructions. Briefly, fibroblasts were seeded in 96-well plates at a density of 0.7 × 10⁴ cells per well in DMEM supplemented with 10% FBS and allowed to adhere overnight. Cells were then treated with serial dilutions of F0565-0303 or vehicle control (DMSO) for 48 h. Following treatment, MTT reagent was added to each well and incubated for 3 h at 37°C. Formazan crystals formed by metabolically active cells were solubilized, and absorbance was measured at 570 nm using a BioTek Cytation 5 multimode plate reader (Agilent, USA).

### CK2α Kinase Activity Assay

CK2α kinase activity was measured using a radiometric filter-binding assay based on incorporation of γ-³³P-ATP (Hartman Analytic, Cat #SCF-3-1-12) into peptide substrate. Recombinant CK2α was pre-incubated with substrate in reaction buffer at room temperature in the presence or absence of test compounds before reaction initiation. The reaction buffer consisted of 20 mM HEPES (pH 7.5), 10 mM MgCl₂, 1 mM EGTA, 0.01% Brij-35, 0.02 mg/mL BSA, 0.1 mM Na₃VO₄, 2 mM DTT, and 1% DMSO. Briefly, CK2tide (Peptide Speciality, Cat#: PSL-CK2tide) as substrate was prepared in freshly made reaction buffer, followed by the addition of CK2α (ProQinase, Cat #0124-0000-1) and gentle mixing. Test compounds (dissolved in 100% DMSO) were dispensed into the reaction mixture using acoustic dispensing technology (Echo 550) at nanoliter volumes and incubated for 20 min at room temperature to allow compound–enzyme interaction. The reaction was initiated by the addition of γ-³³P-ATP and allowed to proceed for 2 h at room temperature. Reactions were terminated by spotting aliquots onto P81 phosphocellulose filter paper, which selectively binds phosphorylated substrate. Unincorporated γ-³³P-ATP was removed by extensive washing with phosphoric acid solution. The retained radioactivity on the filters was quantified as a measure of CK2α kinase activity.

### Bleomycin-induced dermal fibrosis model and F0565-0303 treatment

A murine model of dermal fibrosis was established using bleomycin (MedChemExpress, Cat# HY-17565) as previously described with minor modifications ^(54)^. Briefly, mice were randomly assigned to one of four groups: (i) Vehicle control, (ii) Bleomycin, (iii) Bleomycin + F0565-0303, and (iv) F0565-0303 alone (n=5 per group). Dermal fibrosis was induced by subcutaneous injection of bleomycin sulfate (3.0 mg/kg; MedChemExpress, Cat# ab211091; dissolved in sterile phosphate-buffered saline [PBS]) into the shaved dorsal skin every other day for 14 days. Vehicle control mice received PBS according to the same schedule. Following the initial 14-day fibrosis induction period, mice in the Bleomycin + F0565-0303 group received F0565-0303 at a dose of 15 mg/kg once daily for an additional 14 days. To maintain the fibrotic phenotype throughout the treatment phase, bleomycin administration was continued according to the same dosing schedule. Mice in the F0565-0303 alone group received the compound daily for 14 days in the absence of bleomycin treatment. F0565-0303 was prepared as a 40 mg/mL stock solution in DMSO and diluted immediately before administration in PBS containing 4% Tween-80. Vehicle-treated animals received an equivalent volume of the corresponding DMSO/Tween-80/PBS formulation. At the end of the treatment period (Day 28), animals were euthanized, and dorsal skin tissues from the treatment area were harvested for subsequent histological, biochemical, and molecular analyses. Skin samples were fixed in 10% neutral-buffered formalin for histopathological evaluation.

### Hematoxylin and Eosin (H&E) and Masson’s Trichrome Staining

Paraffin-embedded tissue sections (5 μm) were deparaffinized in xylene (2 changes, 5 min each) and rehydrated through a graded ethanol series consisting of 100% ethanol (2 changes, 3 min each), 95% ethanol (3 min), 80% ethanol (3 min), and 70% ethanol (3 min), followed by rinsing in distilled water. For hematoxylin and eosin (H&E) staining, sections were stained with hematoxylin (StatLab, Cat #SL90-16) for 1–2 min, washed in running tap water for approximately 7 min, and then washed twice in tap water for 10 min each. Sections were then immersed in 80% ethanol for 1 min and counterstained with Eosin Y (Epredia, Cat #71204) for 30 s. Following staining, sections were dehydrated through graded ethanol solutions (70%, 80%, 95%, and 100%), cleared in xylene (2 changes, 5 min each), and mounted with permanent mounting medium. For Masson’s trichrome staining, a commercial kit (Abcam, Cat # ab150686) was used according to the manufacturer’s instructions to assess collagen deposition and tissue fibrosis. Following deparaffinization and rehydration, sections were incubated in preheated Bouin’s solution (60 °C) for 60 min and then washed thoroughly in running tap water. Nuclei were stained with Weigert’s iron hematoxylin for 5 min. Sections were subsequently stained with Biebrich scarlet–acid fuchsin solution for 15 min, differentiated in phosphomolybdic/phosphotungstic acid solution, and stained with aniline blue for 10 min. After brief treatment with 1% acetic acid, sections were dehydrated through graded ethanol solutions, cleared in xylene (2 changes, 5 min each), and mounted with synthetic resin. Collagen fibers stained blue, cytoplasm and muscle fibers stained red, and nuclei stained black. All stained sections were examined and imaged using a brightfield microscope (BZ-X710, KEYENCE, Osaka, Japan) equipped with the manufacturer’s imaging software.

### RNA isolation and bulk RNA sequencing

Total RNA was extracted from tissues using TRIzol Reagent (Thermo Fisher Scientific, Cat #15596026) according to the manufacturer’s protocol. RNA concentration was determined using spectrophotometric methods, and RNA integrity was assessed using an Agilent Bioanalyzer. Only samples with an RNA integrity number (RIN) greater than 9 were used for library preparation. Polyadenylated [poly(A)+] RNA was enriched from 500 ng–1 μg of total RNA using the NEBNext Poly(A) mRNA Magnetic Isolation Module (New England Biolabs, Cat # E7490L) following the manufacturer’s instructions. Briefly, mRNA was captured using oligo(dT) magnetic beads and purified from ribosomal and other non-polyadenylated RNA species. Purified mRNA was fragmented under elevated temperature and used as input for cDNA synthesis. Strand-specific RNA sequencing libraries were prepared using the NEBNext Ultra II Directional RNA Library Prep Kit (New England Biolabs, Cat #7760L). Fragmented mRNA was reverse transcribed to generate first-strand cDNA, followed by second-strand synthesis incorporating dUTP to preserve transcript strand information. Double-stranded cDNA was subjected to end repair, 5′ phosphorylation, dA-tailing, adaptor ligation, and PCR enrichment according to the manufacturer’s recommendations. Library size distribution and quality were evaluated using an Agilent Bioanalyzer, and library concentrations were determined before pooling. Indexed libraries were pooled at equimolar concentrations and sequenced on an Illumina NovaSeq 6000 platform. Raw sequencing reads were demultiplexed and subjected to downstream quality-control and bioinformatic analyses as described below.

### CUT&RUN library preparation and data analysis

CUT&RUN experiments were performed using the CUTANA CUT&RUN Kit (EpiCypher, Cat #14-1001-48s1) according to the manufacturer’s instructions. Briefly, nuclei were isolated from fibroblasts using Wash Buffer (20 mM HEPES, 150 mM NaCl, 0.5 mM spermidine, and 1× Roche cOmplete protease inhibitor cocktail). Nuclei were immobilized on Concanavalin A (ConA) magnetic beads and incubated overnight at 4°C with 0.5 μg of anti-RUNX2 (Cell Signaling, Cat #12556S) antibody in Antibody Buffer. Following antibody incubation, CUTANA pAG-MNase was added and incubated for 10 min at room temperature. Enzymatic digestion was initiated by the addition of CaCl₂ and carried out for 2 h at 4°C before termination with STOP buffer. Samples were subsequently incubated at 37°C for 10 min, and released DNA fragments were purified using the CUTANA DNA Purification Kit according to the manufacturer’s protocol. Sequencing libraries were generated from purified CUT&RUN DNA using the NEBNext Ultra II DNA Library Prep Kit (New England Biolabs, Cat# E7645S). CUT&RUN sequencing data were processed using CUT&RUN Tools (v 2.0) with default parameters.

### Establishment of single-cell RNA libraries

Skin organoids were digested and dissociated into a single-cell suspension. Cells were stained with 7AAD (BD Biosciences, Cat #559925). 7AAD-viable cells were flow sorted using a BD FACSAria flow cytometer. Sorted viable cells were processed using the 10xGenomics 3’OCM kit (10xGenomics, Cat #PN-1000779) following the company’s guide. Briefly, sorted single cell suspensions of 4 samples,10x GEM-X OCM 3’ Gel Bead v4 (10xGenomics, Cat # PN-2001126), and partitioning oil (10xGenomics, Cat # PN-2001213) were loaded into GEM-X 3’ OCM Chip (10x Genomics, Cat # PN-2001099) to capture single cells in nanoliter-scale oil droplets by 10xChromiumX instrument and generate Gel Beads-In-Emulsions (GEMs), and each sample was loaded with one set of beads with a unique list of barcodes. Four samples were barcoded separately but multiplexed in the same GEMs. About 60% loaded cells can be captured in oil droplets, each with a gel bead containing oligos bearing a bead-unique barcode and a 3’ end poly T sequence. Individual cells were lysed in droplets, the released mRNA binds to barcoded Poly T, and the full length of single-strand cDNA from every single nucleus is synthesized by the “template switch” mechanism. GEMs containing single-cell barcoded-cDNA were broken, and all single-cell cDNA libraries were pooled together. Pooled cDNA was cleaned up by DynaBeads MyOne™ Silane beads (10xGenomics, Cat # PN-2000048) and then preamplified by PCR to generate sufficient mass for sequencing library construction. The sequencing libraries were constructed as 3’-biased whole transcriptome libraries by following the steps of cDNA fragmentation, end repair & A-tailing adaptor ligation, and sample index PCR amplification. The final constructed 3’-biased single cell libraries were sequenced by Illumina NextseqX plus machine, targeting 20,000 read pairs/cell, and the sequencing cycles consisted of 28bp for read 1, 90 bp for read 2, and 10 bp for i7 and i5. The sequencing data were processed by the 10xGenomics Cell Ranger pipeline to generate Loupe files or H5 files, which were analyzed by the R Seurat package.

### Single-cell data analysis

scRNA-seq data from three samples (Control, Day-8, and Day-14) were analyzed in R (version 4.2-4.3 series) using Seurat v5. Raw count matrices were generated using 10x Genomic platform and were imported with Read10X(), and Seurat objects were created for each sample using CreateSeuratObject() with min.cell = 3 and min.features = 200. We subsampled 2,000 cells per sample to ensure the balanced representation. Cells were integrated based on the top 2000 differentially expressed (DE) genes in the combined dataset. PCA analysis was performed retaining 30 principal components. The first 20 principal components were used for making shared nearest-neighbor graph construction using the Louvin algorithm at resolution of 0.5 and UMAP was created for visualization. This yielded a total of 18 clusters. Markers genes were identified by using FindAllMarkers() with the Wilcoxon rank-sum test (only.pos = TRUE, min.pct = 0.25, logfc.threshold = 0.25), and pairwise differential expression analyses between Control and treatment groups were performed using FindMarkers(). Using canonical genes from the literature, we labelled each cluster with its cell type. Gene expression pattern were visualized using FeaturePlot() and VlnPlot(). Subsequent visualization was performed using R packages like dplyr(v1.2.0), ggplot2 (v4.0.2) and patchwork (v1.3.2).

### Trajectory inference and pseudotime analysis

For trajectory inference, we used Slingshot (version) with cluster labels as input and the UMAP embedding as the reduced-dimensional space. Cluster 12 was specified as the root cluster based on its biology and position in the UMAP manifold. Slingshot (v2.16.0) was applied to the full dataset without subsetting to infer a global trajectory across all samples. Pseudotime values from the merged Lineage-5-8 trajectory were used as continuous measures of cellular progression. Psuedotime values were mapped onto the shared UMAP embedding, and sample-wise distributions were compared using density plots.

Cells with defined pseudotime values were retained, and coordinates were extracted from the UMAP. Local directional vectors were estimated in the two-dimensional UMAP space by performing k-nearest neighbour search (k = 25) and identifying neighbouring cells with higher pseudotime values. The mean displacement vector toward higher-pseudotime neighbors were calculated and normalized to unit length for each cells and these per-cell vectors were then aggregated on a two-dimensional grid spanning the UMAP space, grid bins having fewer than 4 cells were excluded. The bins with weaker directionality below the 10^th^ percentile of vector length were removed. The surviving vectors were rescaled to a constant arrow length so the arrow only denotes the direction not the magnitude. Sample-specific vector fields were overlaid on UMAP plots, with cells colored by Lineage-5-8 to visualize relative progression along the inferred trajectory in each condition providing a pseudotime-gradient-based visualization.

Pseudotime heatmap was generated using the pseudotime-dependent gene expression along the merged Lineage-5-8. Normalized log-expression values from the RNA assay were extracted and for each gene Spearman correlation was computed with Lineage-5-8 pseudotime vector. Adjusted p-values were retrieved by applying Benjamini-Hoshberg correction and genes were ranked by significance and correlation sign which was used for ranking the genes. GO analysis was performed intersecting the GO gene sets for with our genes in our datasets.

### Sample-level pseudotime statistics

To quantify differences in cellular progression across samples, we analyzed the distribution of cells along the merged Lineage-5-8 pseudotime trajectory using sample-level summaries as the primary units of comparison. Cells belonging to the trajectory of interest were defined as those with non-missing Lineage-5-8 values. Continuous pseudotime values were ranked across the pooled sets of cells from all samples and partitioned into three equally sized bins, which were selected for categorical comparisons for Early, Middle, and Late pseudotime states. For donor-level comparisons, the proportions of cells in each pseudotime bin were calculated separately for Control, Day 8, and Day 14. To examine patient-specific deviations from the control as

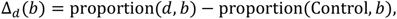

where *b* ∈ {Early, Middle, Late}. These differences were visualized using line and point plots to compare cell redistribution along pseudotime among donors.

For exploratory statistical assessment of enrichment or depletion, Fisher’s exact tests were performed separately for each donor pair (Control vs Day 8 and Control vs Day 14). For a given bin, cells were classified as either belonging to that bin or not, and 2 × 2 contingency tables were constructed from the corresponding cell counts in the disease and control samples. Fisher’s exact test was then used to estimate odds ratios and P values for bin-specific enrichment or depletion in each disease donor relative to the control. Because these tests treat cells as independent observations and do not fully account for donor-level dependence, they were interpreted as exploratory cell-level measures rather than definitive inferential statistics.

### PROGENy (v1.30.0) pathway activity heatmap analysis

Pathway activity analysis was performed to summarize signaling programs across selected fibroblast-associated clusters in Control, Day 8, and Day 14 samples. To estimate pathway activity at the sample-by-cluster level, a grouped average expression matrix was generated from normalized RNA expression values. Group-wise mean expression values were then calculated for each gene across all sample-cluster combinations, producing a gene-by-group expression matrix. Pathway activity scores were computed using the human PROGENy (v1.30.0) model with the top 500 footprint genes per pathway. The intersection between genes present in the expression matrix and genes represented in the PROGENy weight matrix was used to define the final input gene set. For each sample-cluster group, pathway activity was calculated as a weighted sum of expression values across footprint genes, resulting in a pathways-by-group score matrix. PROGENy (v.1.30.0) uses signed pathway weights; resulting scores reflect relative pathway activation or repression across groups.

The analysis focused on fibroblast-related and neighbouring clusters. Accordingly, pathway scores were restricted to cluster 0, 1, 5, 6, 9, 12, and 13. To simplify downstream visualization, clusters 1 and 13 were merged at the pathway-score level by averaging their values within each sample, generating a composite cluster labeled 1. After merging, the retained set of clusters was {0, 1, 5, 6, 9, 12}. Pathway scores were standardized across sample-cluster groups by row-wise z-score scaling so that each pathway had a mean of zero and unit variance across the displayed columns. Heatmap was generated using pheatmap (1.0.13).

### Cell-cell communication analysis

Cell-cell communication analysis was performed using CellChat (v1.6.1). The dataset contained approximately 6,000 cells, and 40,606 and RNA assay was used for all CellChat analyses. Normalized expression values from the RNA assay were used to infer ligand-receptor communication. Cells were already annotated with sample identity (Control, Day 8, and Day 14), Seurat (v5) cluster assignments, and curated cell-type labels.

Two strategies were to define communicating cell populations. For fibroblast-keratinocyte communication analysis, cells were grouped into Fibroblast, Keratinocyte, and Other categories on the basis of cluster identity. For fibroblast subset communication analysis, fibroblast populations were further subdivided into Fibro_Sender_1_5 (Seurat clusters 1 and 5) and Fibro_Receiver_0_6_9 (Seurat clusters 0, 6, and 9), while keratinocyte and other populations were retained to preserve the surrounding cellular context. This second grouping enabled directional analysis of fibroblast subset signaling. CellChat (v1.6.1) was run on normalized RNA expression data using the human ligand-receptor reference database (CellChatDB.human) with default parameters. Communication probabilities were inferred and interactions involving fewer than 10 cells per group were excluded.

To evaluate the directional signaling, the following communication modes were examined separately in each sample, 1: Fibroblast to Keratinocyte and Fibro_Sender_1_5 to Fibro_Receiver_0_6_9. For each sample and communication mode, signaling pathways were ranked according to total communication probability.

### Dataset integration and batch correction using Harmony (v1.2.4)

We downloaded the publicly available datasets (write names of the datasets), and 19,683 cells were taken from this set, and this set was compared to our lab dataset containing 6000 cells. The publicly available dataset was preprocessed, including normalization, identification of 3,000 highly variable genes using the variance-stabilizing transformation method, scaling, and principal component analysis (PCA) with 30 principal components. The PCA embedding was extracted from the Seurat object and aligned with cell-level metadata for integration.

Batch correction was performed using Harmony (v1.2.4) with theta = 8, nclust = 50, and max.iter.harmony = 20. In this context, theta controls the strength of batch correction, with higher values imposing stronger dataset mixing in the corrected embedding; nclust defines the number of soft clusters used internally by Harmony during iterative correction; and max. iter.harmony sets the maximum number of Harmony optimization rounds. The resulting Harmony-corrected matrix was added back to the Seurat object as a new dimension for downstream analysis.

All downstream analyses were performed in the Harmony-corrected space. A two-dimensional UMAP embedding was generated using the first 30 Harmony dimensions with n.neighbors = 50 and min.dist = 0.5. The increased n.neighbors parameter was used to emphasize broader manifold structure and improve cross-dataset continuity, whereas the increased min.dist value produced a more diffuse layer that facilitated visualization of dataset overlap with integration. Shared nearest-neighbor graph construction and Louvain clustering were then performed using the first 30 Harmony dimensions. Cluster assignments were stored in Seurat_clusters, and UMAP visualization colored by dataset and cluster identity was used to assess integration quality and cluster structure. The integrated Seurat object, including Harmony embeddings, UMAP coordinates, graph structure, and cluster assignments, was saved for downstream analysis.

### Bulk RNA-seq data preprocessing, alignment, and quantification

Raw paired-end bulk RNA-seq FASTQ files were processed on the UAB Research Computing platform Cheaha. Adapter removal and quality trimming were performed with Trimmomatic v0.36 in paired-end mode using the TruSeq3-PE adapter set and the following parameters: ILLUMINACLIP:2:30:10, LEADING:3, TRAILING:3, SLIDINGWINDOW:4:15, and MINLEN:36. Trimmed paired-end reads were then aligned to the human reference genome (hg38) using STAR v2.7.3a on the Cheaha cluster. Alignments were output as coordinate-sorted BAM files. For downstream gene-level quantification, aligned reads were counted HTSeq-count v2.0.3 with the hg38 reference annotation file (hg38.refGene.gtf). Counting was performed on coordinate-sorted BAM files with parameters specifying BAM input format, position-sorted reads, reverse-strand specificity, exon-level feature assignment, gene_id summarization, union-mode overlap handling, and a minimum alignment quality score of 10 (-f bam -r pos -s reverse -t exon - i gene_id -m union -a 10).

Differential expression analysis was performed in R using DESeq2 v1.48.2 on gene-level count matrices generated with HTSeq-count v2.0.3. The TGF-specific RUNX2 knockdown effect was obtained from the contrast representing shRUNX2 versus shCtrl under TGF treatment. Genes with padj < 0.10 and log2FoldChange < 0 were defined as the loose TGF-downregulated set. Volcano plots were used to visualize differential expression results, with significance defined as padj < 0.05 and |log2FoldChange| ≥ 1 unless specified differently.

### Gene Ontology analysis

GO enrichment analysis was performed in R using clusterProfiler (v4.16.0) with the human annotation database org.Hs.eg.db (v3.21.0). Enrichment analysis focused on Gene Ontology Biological Process terms and was carried out using clusterProfiler::enrichGO(). Enrichment results were summarized in per-condition tables and visualized using bubble plots generated in base R. GO based results were also used to generate gene-concept network plot using enrichplot (v1.28.4).

### Statistical analysis

All statistical analyses were performed using GraphPad Prism 11 (GraphPad Software, La Jolla, CA, USA) or by the default algorithms implemented in R- or Python-based analysis pipelines. Unless otherwise specified, data are presented as mean ± standard deviation (SD). Statistical significance between groups was assessed using one-way analysis of variance (ANOVA), Wilcoxon rank-sum test, or Fisher’s exact test, as appropriate. Multiple independent experiments were conducted to confirm the reproducibility of all findings. A *P* value < 0.05 was considered statistically significant unless otherwise stated.

---

**Figure S1.**
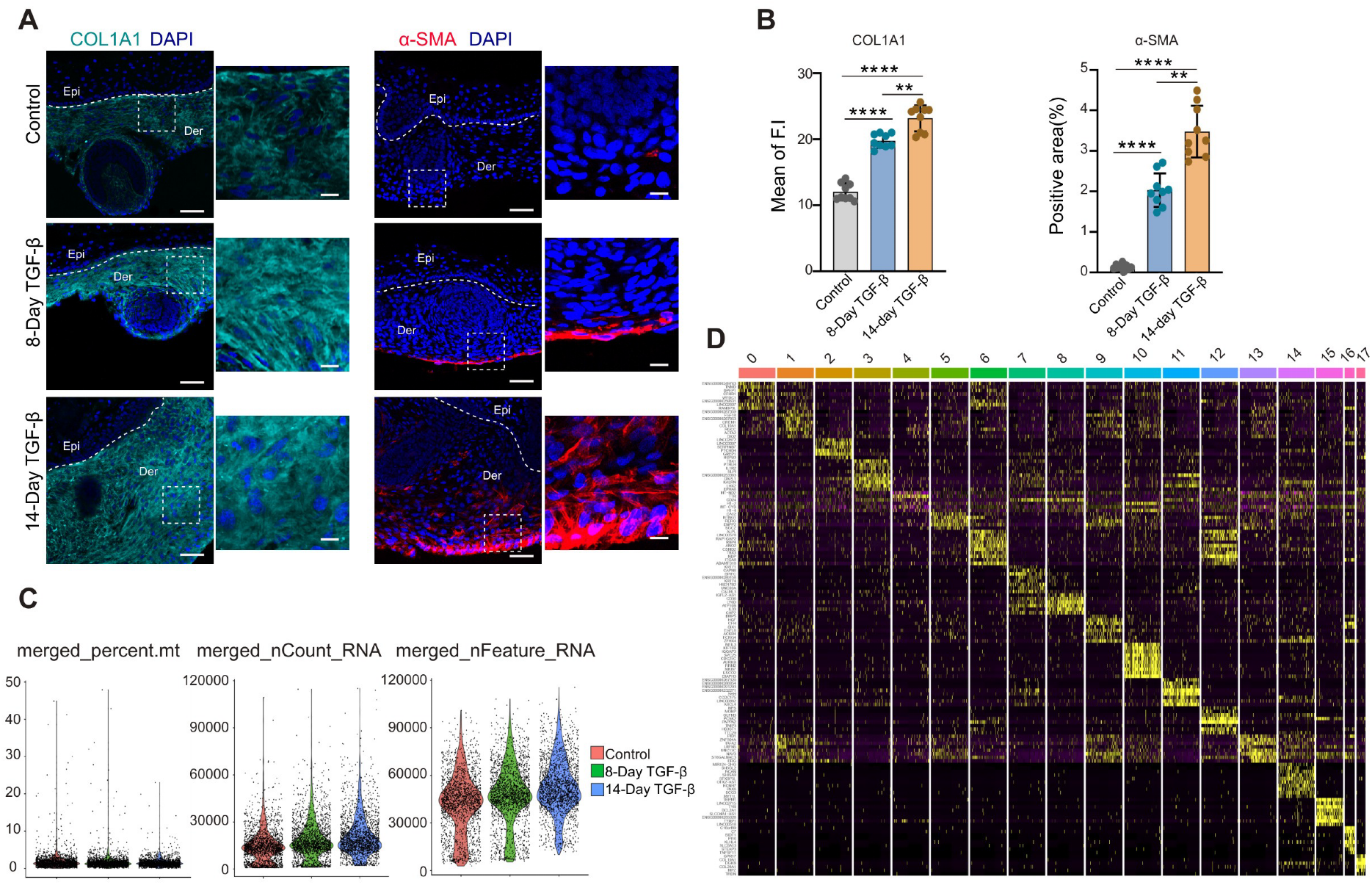
Single-cell RNA sequencing reveals the quality and cellular composition of TGF-β–treated skin organoid datasets. (A, B) Immunofluorescence (IF) analysis demonstrates increased COL1A1 and α-SMA expression in an independent iPSC-derived skin organoids following 8- and 14-day TGF-β treatment, with corresponding quantification of fluorescence intensity. For each condition, n = 3 independent organoids were analyzed, with 3 randomly selected regions per organoid. Epi, epidermis; Der, dermis. White dot line: boundary between epidermis and dermis. Scale bar, 100 μm. Scale bar, 100 μm. (B) Quality control metrics showing distribution of detected gene numbers (nFeature_RNA) and total transcript counts (nCount_RNA) per cell across samples. The percentage of mitochondrial gene expression across cells is used to filter out low-quality or stressed cells. Cells with high mitochondrial content were excluded before downstream analysis. (D) heatmap showing the top differentially expressed genes across identified cell clusters, illustrating cluster-specific transcriptional signatures used for cell-type annotation. Data in (B) is presented as mean ± SEM. One-way ANOVA determined statistical significance with multiple comparisons (\*\**P* < 0.01, \*\*\*\**P*<0.0001).

**Figure S2.**
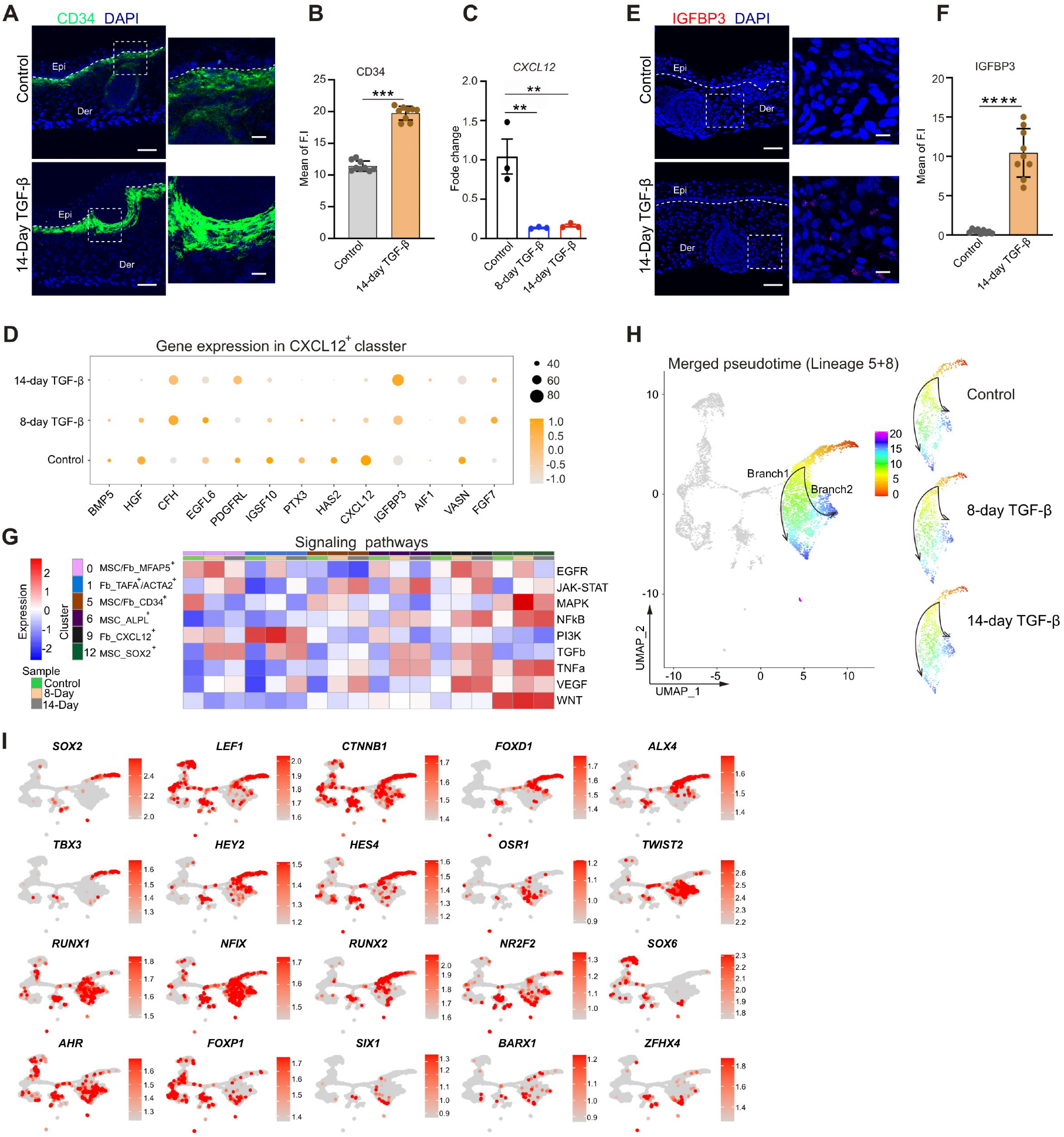
Pseudotime trajectory reveals bifurcation of MSC/Fb-MFAP5⁺ cells and associated transcription factor programs. (A, B) IF staining of CD34 in control and TGF-β–treated independent iPSC-derived skin organoids following 14-day treatment, with corresponding quantification of fluorescence intensity. For each condition, n = 3 independent organoids were analyzed, with 3 randomly selected regions per organoid. Scale bar, 100 μm. (C) qPCR analysis of *CXCL12* expression in control, 8-day, and 14-day TGF-β–treated organoids. (D) Bubble plot summarizing CXCL12^+^ cluster-specific transcriptional responses. (E, F) IF staining of IGFBP3 in control and TGF-β–treated independent iPSC-derived skin organoids following 14-day treatment, with corresponding quantification of fluorescence intensity. For each condition, n = 3 independent organoids were analyzed, with 3 randomly selected regions per organoid. Epi, epidermis; Der, dermis. White dot line: boundary between epidermis and dermis. Scale bar, 100 μm. Scale bar, 100 μm. (G) Heatmap showing alterations in the indicated signaling pathways in control, 8-day, and 14-day TGF-treated organoids. (H) Pseudotime trajectory analysis of dermal populations projected onto the integrated UMAP of control and TGF-β–treated organoids, revealing two branching trajectories originating from MFAP⁺ cells, with one branch toward TAFA^+^/ACTA2⁺ cells and the other toward CD34⁺ cells. (I) Feature plots showing the expression of representative transcription factor (TF) genes identified from pseudotime analysis across dermal cell populations. Data in (B), (C), and (F) are presented as mean ± SEM. One-way ANOVA determined statistical significance with multiple comparisons (\*\**P* < 0.01, \*\*\**P* < 0.001, ****P < 0.0001).

**Figure S3.**
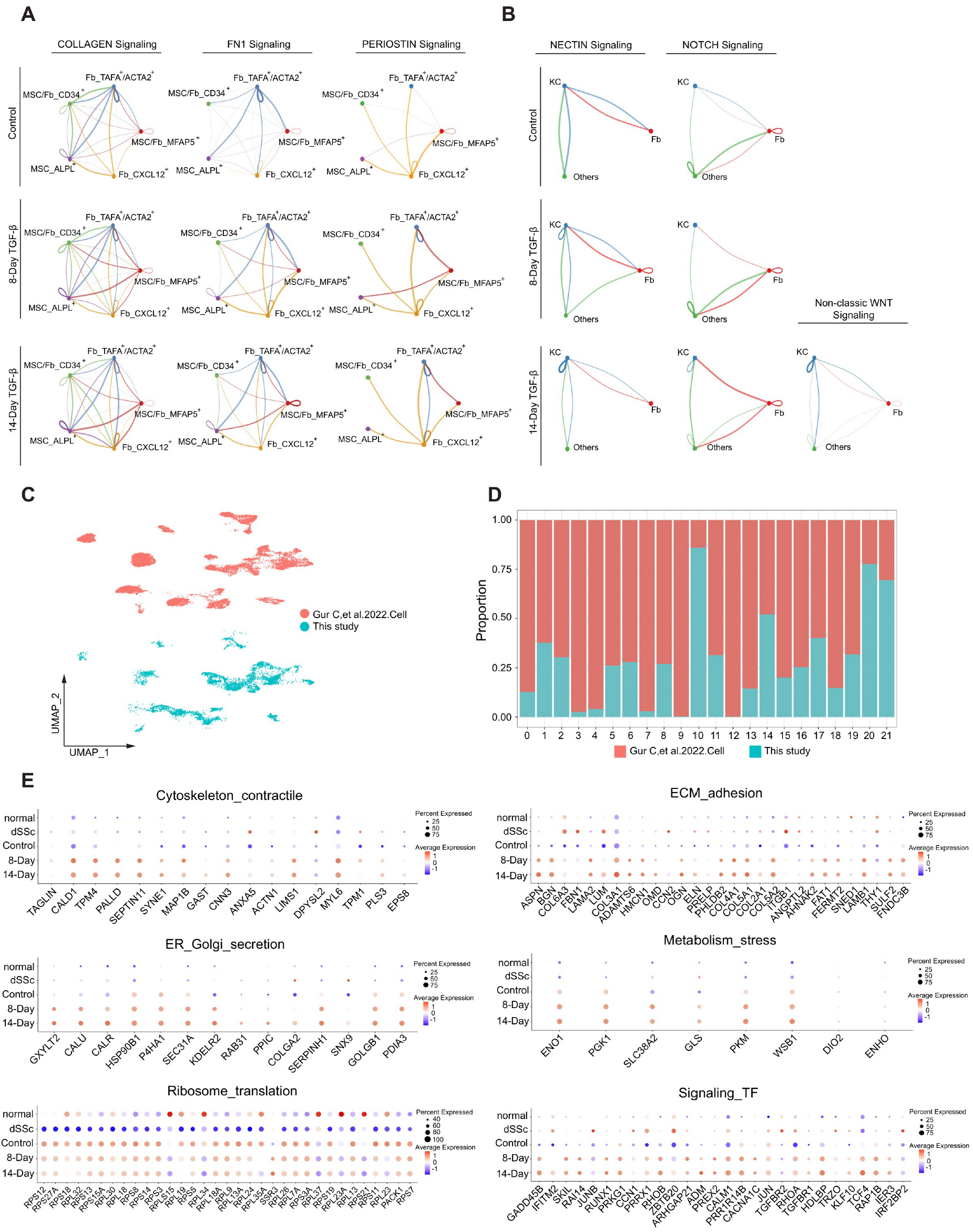
Cellular crosstalk and gene regulatory changes in fibrotic organoids parallel those found in human scleroderma skin. (A, B) CellChat analysis of subfibroblasts (A) and keratinocyte-fibroblast (B) communication networks in control and TGF-β–treated organoids at 8-day and 14-day time points, showing the relative contribution and distribution of top-ranked signaling pathways across both cell types. (C, D) UMAP visualization of cell population distribution (C) and proportion of cells in each subset (D) from fibrotic organoid and human scleroderma skin datasets, enabling cross-system comparison of cellular composition. (E) Bubble plot comparing pseudo-bulk expression levels of genes within the indicated functional categories between human scleroderma and fibrotic organoid datasets.

**Figure S4:**
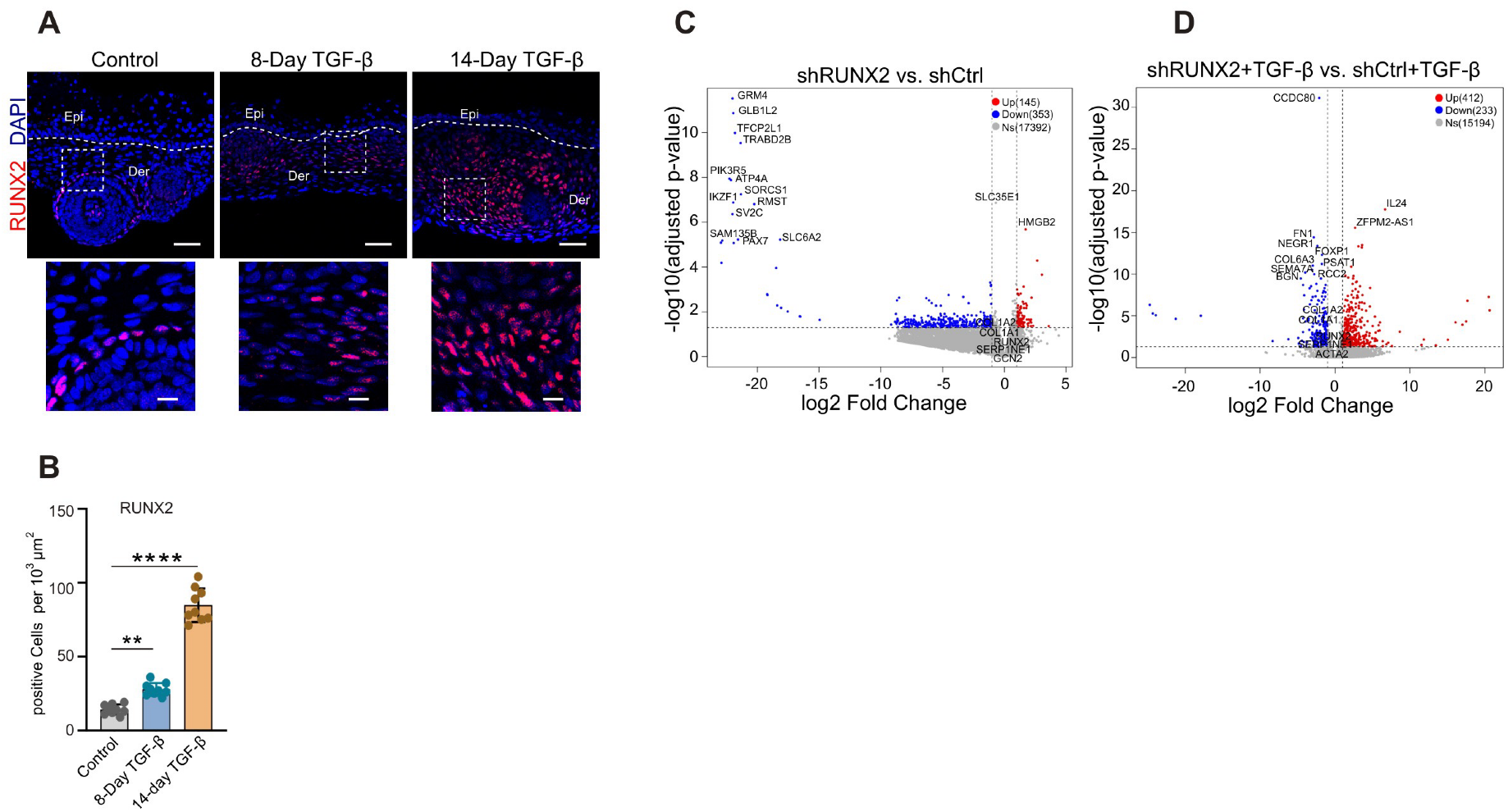
RUNX2 knockdown alters transcriptional programs under basal and TGF-β conditions. (A, B) IF staining of RUNX2 in control and TGF-β–treated independent iPSC-derived skin organoids at 8-day and 14-day time points, with corresponding quantification of fluorescence intensity. For each condition, n = 3 independent organoids were analyzed, with 3 randomly selected regions per organoid. Epi, epidermis; Der, dermis. White dot line: boundary between epidermis and dermis. Scale bar, 100 μm. Scale bar, 100 μm. (C, D) Volcano plots showing differential gene expression between shCtrl and shRUNX cells under basal and TGF-treated conditions, with top differentially expressed genes highlighted and labeled. Data in (B) is presented as mean ± SEM. One-way ANOVA determined statistical significance with multiple comparisons (\*\**P* < 0.01, ****P < 0.0001).

**Figure S5:**
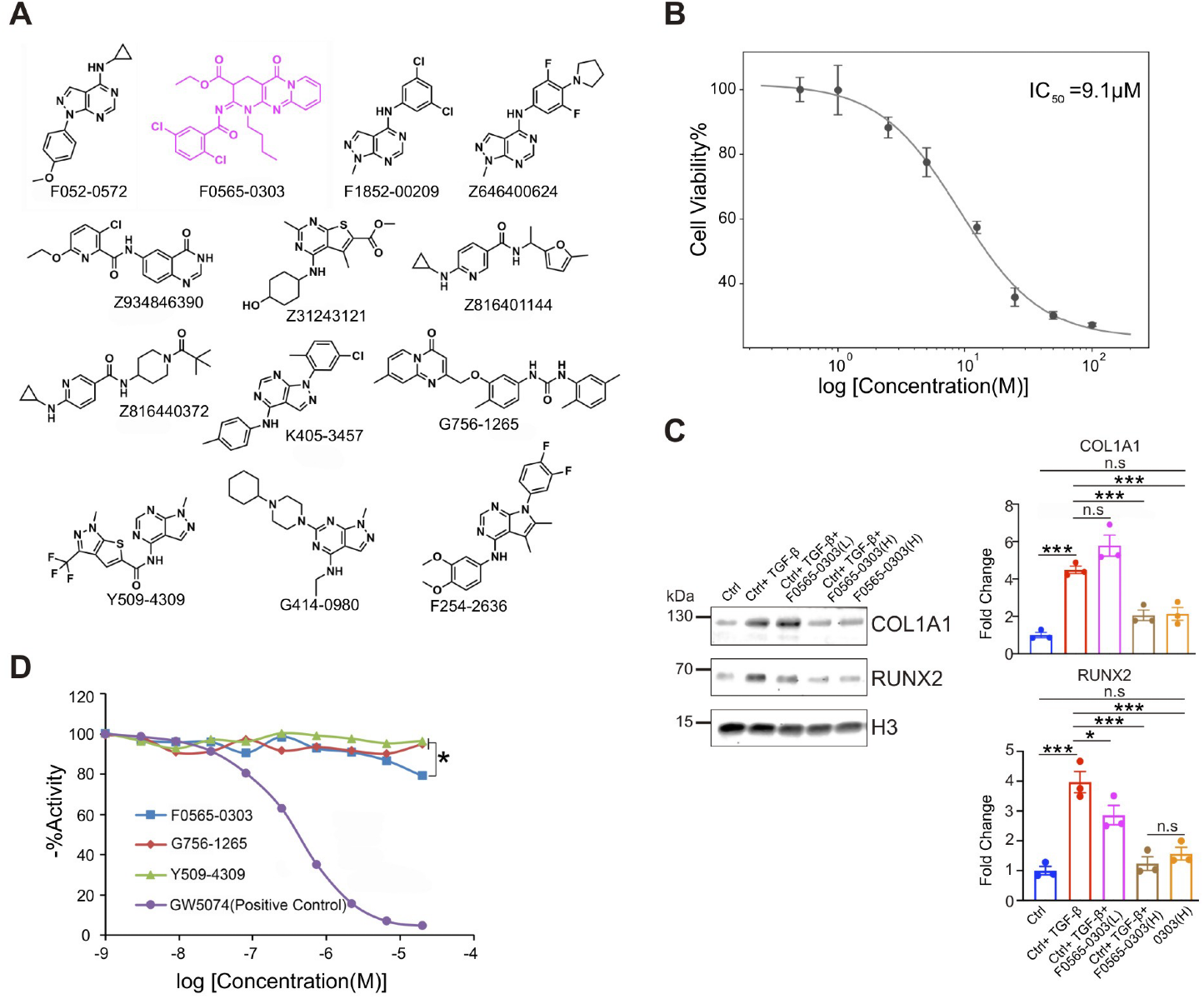
Pharmacological analyses demonstrate the potent anti-fibrotic effects of F0565-0303. (A) Chemical structures of 13 candidate compounds used for initial screening. (B) Determination of the IC50 value of F0565-0303 based on the cell viability analysis. (C) Western blot analysis shows RUNX2, COL1A1, and α-SMA protein expression in Pr-Fbs treated with F0565-0303 at 0.5 μM and 5 μM for 48 hours in the presence or absence of TGF-β. (D) CK2 kinase activity assay evaluating the inhibitory effect of F0565-0303, G756-1265, and Y504-4309 on CK2α enzymatic activity *in vitro*. GW5074, cRaf1 kinase inhibitor, served as a positive control. One-way ANOVA determined statistical significance with multiple comparisons (\**P* < 0.05).

**Figure S6:**
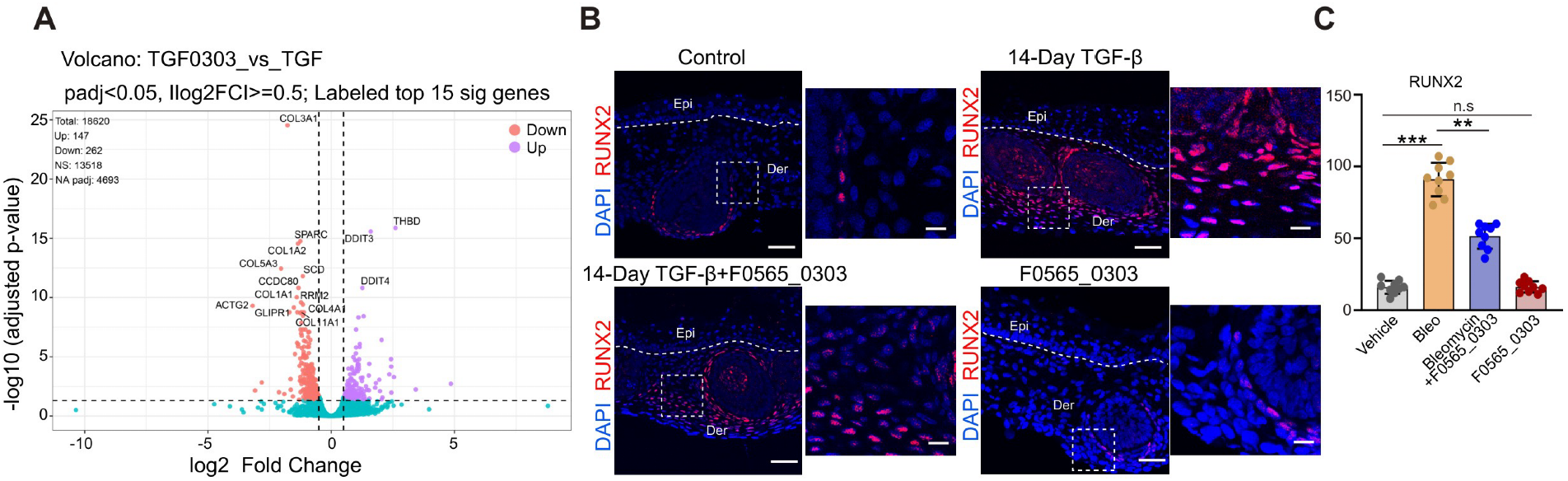
F0565-0303 modulates fibrotic gene expression and dermal fibrosis across cellular and *in vivo* models. (A) Volcano plots showing differential gene expression in Pr-Fbs under F0565-0303-treated conditions, with top differentially expressed genes highlighted and labeled. (B, C) IF staining of RUNX2 in an independent iPSC-derived skin organoids treated with TGF-β for 14 days, with 5 μM F0565-0303 co-treatment administered during the final 8 days of the treatment period. Quantification of fluorescence intensity is shown. For each condition, n = 3 independent organoids were analyzed, with 3 randomly selected regions per organoid. Epi, epidermis; Der, dermis. White dot line: boundary between epidermis and dermis. Scale bar, 100 μm. Scale bar, 100 μm. Data in (C) is presented as mean ± SEM. One-way ANOVA determined statistical significance with multiple comparisons (\*\**P* < 0.01, \*\*\**P* < 0.001).

